# Layer 1 of somatosensory cortex: An important site for input to a tiny cortical compartment

**DOI:** 10.1101/2021.11.26.469979

**Authors:** Julia MT Ledderose, Timothy A Zolnik, Maria Toumazou, Thorsten Trimbuch, Christian Rosenmund, Britta J Eickholt, Dieter Jaeger, Matthew E Larkum, Robert NS Sachdev

## Abstract

Neocortical Layer (L) 1 has been proposed to be at the center for top-down and bottom-up integration. It is a locus for interactions between long-range inputs, L1 interneurons and apical tuft dendrites of pyramidal neurons. While input to L1 has been studied intensively, the level and effect of input to this layer has still not been completely characterized. Here we examined the input to L1 of mouse somatosensory cortex with retrograde tracing and optogenetics. Our assays reveal that local input to L1 is predominantly from L2/3 and L5 pyramidal neurons and interneurons, and that subtypes of local L5 and L6b neurons project to L1 with different probabilities. Long-range input from sensory-motor cortices to L1 of S1 arose predominantly from L2/3 neurons. Our optogenetic experiments showed that intra-telencephalic L5 pyramidal neurons drive L1 interneurons but have no effect locally on L5 apical tuft dendrites. Dual retrograde tracing revealed that a fraction of local and long-range neurons were both presynaptic to L5 neurons and projected to L1. Our work highlights the prominent role of local inputs to L1 and shows the potential for complex interactions between long-range and local inputs which are both in position to modify the output of somatosensory cortex.

## Introduction

Neocortical layer (L) 1, long an enigma (Burkhalter 1989; Gămănuţ et al. 2018; Hubel 1982; Ibrahim et al. 2020; Marín-Padilla 1998), has recently become central to ideas about consciousness and perception (Aru et al. 2020; Gidon et al. 2020; Guest et al. 2021; Ibrahim et al. 2021; M. Larkum 2013; Schroeder et al. 2022; Suzuki and Larkum 2020; Takahashi et al. 2016; Takahashi et al. 2020). These ideas seem astonishing for such a cell sparse, tiny layer, located at the surface of cortex, spanning only 100 microns in width in mice, and containing only GABAergic interneurons (Rudy et al. 2011; Schuman et al. 2021). While L1 is sparse in cell bodies, it is filled with processes, both dendrites of a variety of local pyramidal neurons and interneurons, and axons of a variety of inputs (L. Cauller 1995; DeFelipe and Fariñas 1992; DeFelipe 1997; Gabbott and Somogyi 1986; Ibrahim et al. 2020; Ito et al. 1998; Karimi et al. 2020; M. Larkum 2013; Lee et al. 2010; Marín-Padilla 1998; Mason and Larkman 1990; Schuman et al. 2019). A consistent theme for L1 in all cortices is that L2/3 and L5 pyramidal neurons have dendrites in L1, which makes this layer a potential locus for contextual interactions between the dendrites of the pyramidal neurons and long-range feedback inputs (Abs et al. 2018; Anastasiades et al. 2021; Cruikshank et al. 2012; Doron et al. 2020; Guest et al. 2021; Rudy et al. 2011; Schuman et al. 2021). This organization of inputs and apical tuft dendrites has led to the proposal that the apical tuft dendrites in L1 are a key element for sensory perception and learning.

One long standing related idea is that primary sensory cortical areas generate feedforward input to higher order cortical areas, to the middle layers of cortex, which in turn provide feedback to lower order cortical areas, to the outer layers - including L1 - of cortex (Felleman and Van Essen 1991; Markov et al. 2014; Rockland and Pandya 1979). This work suggested specific laminar profiles of cortico-cortical connectivity with a class of long-range inputs arising exclusively from infragranular layers, and a second class from both infragranular and supragranular layers (Felleman and Van Essen 1991). These ideas have been synthesized into a hypothesis with the pyramidal neuron and its apical tuft dendrites as a key locus for the contextual interaction between feedforward and feedback input (M. Larkum 2013). According to this hypothesis, when a pyramidal neuron receives strong feedforward input targeting its somatic region, action potentials are triggered that backpropagate into the apical dendrite; if feedback inputs activate the apical tufts at the same time (Takahashi et al. 2020), this leads to long lasting calcium potentials in the apical dendrite and bursts of somatic action potentials (Doron et al. 2020; Guest et al. 2021).

Earlier work with the retrograde tracers fast blue and diamidino yellow has shown that long-range cortico-cortical projections are an important source of input to L1 of somatosensory cortex (L. Cauller 1995; L. J. Cauller et al. 1998). Single-cell fills of motor cortical and thalamocortical neurons also reveal that long-range axons target L1 (Ohno et al. 2012; Rubio-Garrido et al. 2009; Veinante and Deschênes 2003). Quantification of axons and boutons in L1 suggests a key role for cortico-cortical input to this layer (Binzegger et al. 2004; Boucsein et al. 2011; Douglas and Martin 2007). Additionally, retrograde tracing with rabies virus shows that L1 neuron-derived neurotropic factor (NDNF) positive interneurons receive long-range input (Abs et al. 2018; Anastasiades et al. 2021; Cohen-Kashi Malina et al. 2021). This earlier work also shows that a variety of local inputs target L1: Single-cell fills show that axons of local L2/3 and L5 neurons arborize in L1 (Brown and Hestrin 2009; Feldmeyer et al. 2006; Sakmann 2017; Schuman et al. 2021); fast blue application on the cortical surface shows that local L5 and L6b neurons connect to L1 (Clancy and Cauller 1999); paired recordings from L1 interneurons and L2/3 pyramidal cells show that L2/3 neurons can modulate the activity of L1 interneurons (Wozny and Williams 2011); and retrograde tracing with rabies shows that the bulk of input to the NDNF interneurons is from local sources (Abs et al. 2018).

While a lot is known about the input to L1, the earlier work with tracers and single-cell fills did not quantify the extent, the laminar profile or identity of local and long-range inputs to L1 project to L1 locally or from long-range sites. Even the work with rabies, targeting the input to NDNF-Cre positive cells also only reveals input to one class of interneurons, which may not be representative of the overall input to L1. Here we used retrograde labelling with fast blue, retrobeads and rabies, and quantified local and long-range connectivity. We focus on L5 neurons because L5 pyramidal neurons are the main output neurons of cortex, connecting cortex to a variety of subcortical structures -- thalamus, basal ganglia, pons, and spinal cord. Additionally, these neurons extend their apical tuft dendrites into L1 and have an axon that contributes to local recurrent excitation, with local branches that can extend into L1 (Larkum 2013; Sakmann 2017).

Our work indicates that the majority of input to L1 is local, and that the long-range input to L1 of S1 is from a mix of supragranular and infragranular neurons distributed in sensory motor cortices. We identified classes of local and long-range neurons that project to L1, and used circuit mapping to reveal the effect of intratelencephalic neurons on L1 interneurons.

## Methods

### Ethics statement

All experiments were conducted under the license G0278/16 in accordance with the guidelines of animal welfare of the Charité Universitätsmedizin Berlin and the local authorities, the ’Landesamt für Gesundheit und Soziales’.

### Mice

We used adult wildtype C57BL6/J mice and transgenic Ai9 reporter mouse lines expressing layer-specific genes under the tdTomato (tdTom) promoter (Breeding Unit Steglitz, Charité Universitätsmedizin, Berlin): Tlx3-Cre for L5 intracortical (IT) projections (Tlx3-Cre Tg(Tlx3-cre) PL56Gsat/Mmucd (NIMH) MMRRC Stock 041158-UCD, lfd nr. 1287) (Gerfen et al. 2013); Sim1-Cre for L5 pyramidal tract (PT) projections (Sim1-Cre Tg(Sim1-cre) KJ18Gsat/Mmucd, (NIMH) MMRRC Stock 031742-UCD, lfd nr. 1288) (Gerfen et al. 2013), Drd1a-Cre and Ctgf-2A-dgCre for L6b neuronal subpopulations (Drd1a-Cre Tg(Drd1-cre) FK164Gsat/Mmucd, MMRRC Stock 030781-UCD, lfd nr. 1286, Benjamin Judkewitz, Charité; Ctgf-2A-dgCre Cg-Ccn2< tm1.1 (folA/cre) Hze > /J, JAX Stock 028535, Benjamin Judkewitz, Charité), Sst-IRES-Cre for somatostatin positive interneurons (SST-IRES-Cre, Sst<tm2.1(cre)Zjh>/J, JAX Stock 013044), Vip-IRES-Cre for vasoactive intestinal peptide (VIP-IRES-Cre, Vip<tm1(cre)Zjh>/J, JAX Stock 010908), PV-IRES-Cre for parvalbumin positive interneurons (PV-IRES-Cre, Pvalb<tm1(cre)Arbr>/J, JAX Stock 017320); Scnn1a-Cre for layer 4 neurons (Scnn1a-Tg3-Cre, B6;C3-Tg(Scnn1a-cre)3Aibs/J, IMSR_JAX:00961); Gpr26-Cre Tg (Gpr26- cre), lfd nr. 1333, KO250Gsat/Mmucd; MMRRC Stock 033032-UCD (Ehud Ahissar, Weizmann Institute) for neurons of the posteromedial complex of thalamus (POm). We crossed the Cre-lines with Ai9 reporter mice to induce tdTom expression in Tlx3, Sim1, Drd1, Ctgf, Scnn1a, Vip, Sst, and PV neurons. The Cre expression in Ctgf-Cre mice was stabilized by intraperitoneal administration of trimethoprim (250 µg/g body weight). All mice were kept in groups of two to three individuals under standard conditions in a 12-hour day-night cycle at 21˚C room temperature. Water and food were available ad libitum.

### Rabies virus production

All viruses for rabies tracing were produced in the Viral Core Facility of the Charité, Universitätsmedizin Berlin (vcf.charite.de). The production was previously described by the Callaway lab (Osakada and Callaway 2013). In brief, we transfected B7GG cells by Lipofectamine 3000 (Thermo Fisher) with the rabies virus genomic vectors that contained either mCherry (pSADdeltaG-F3-mcherry, Addgene plasmid #32634), or GFP and Synaptophysin-RFP cDNA (pRVdG-N-P-M-EGFP-SynPhRFP-L, Addgene plasmid #52483), and with these additional plasmids: pcDNA-SADB19N Addgene plasmid #32630, pcDNA-SADB19P Addgene plasmid #32631, pcDNA-SADB19L Addgene plasmid #32632, pcDNA-SADB19G Addgene plasmid #32633. We collected the supernatant over several days and re-transfected the recovered virus in B7GG cells for a final collection step. For additional pseudotyping, the rabies with the envelope protein EnvA of the Avian Sarcoma and Leukosis virus (ASLV), and the obtained supernatant containing unpseudotyped viruses was applied onto BHK-EnvA cells. After three to five days, EnvA-pseudotyped rabies viruses containing supernatant were collected, filtered and concentrated by ultracentrifugation. Rabies virus titer was determined by infecting a serial dilution of the virus in HEK293T-TVA cells. All components for rabies virus production were gifts from Edward Callaway (Salk Institute, San Diego, CA, USA).

To allow expression in Cre positive starter cells, we applied a previously described AAV vector that contained a Cre dependent expression cassette of a nuclear GFP, EnvA interacting cognate avian viral TVA receptor and an optimized rabies G protein (pAAV-Syn-Flex-nGToG-WPRE3) (Choi and Callaway 2011; Kim et al. 2016; Sun et al. 2014; Zolnik et al. 2020). To distinguish between rabies GFP expression and GFP expression from AAV infected starter cells, we exchanged the nuclear GFP with mCerulean3 (addgene plasmid #54730) (Markwardt et al. 2011) to form pAAV-Syn-Flex-Ce3ToG-WPRE3. After sequence verification, AAV was produced at the Viral Core Facility of the Charité – Universitätsmedizin Berlin.

### Preparation of mice for surgeries

We deeply anaesthetized the mice (n=42) according to their weight with a mixture of ketamine (100 mg/kg) and xylazine (10 mg/kg). After full anesthesia, confirmed by toe pinch for the absence of reflexes, mice were placed into a stereotaxic frame with non-puncture ear bars and a nose clamp (Kopf Stereotaxic device, California, USA, Inc.). For analgesics, we applied 100 µl of lidocaine under the scalp before surgery, and administered Carprofen (5 mg/kg) and Buprenorphine (5 mg/kg) intraperitoneal for analgesic post-treatment after surgery. Body temperature was maintained at 37 °C, using a heating pad during the surgery and while mice recovered from the operation.

### Fast blue application

Fast blue (fb) was applied on L1 of S1 cortex, where it is taken up for retrograde transport along the axons (Keizer et al. 1983; Kuypers et al. 1980). We made a craniotomy of ∼ 0.5 mm^2^ over S1 cortex (AP: -1.2 to -1.7 mm, lateral: 2.7 mm to 3.3 mm) and without removing the dura mater applied 2 µl (1% in H_2_O) of the retrograde, blue fluorescent neuronal tracer fast blue (Polyscience Inc, 17740-1, 1mg) onto the surface of L1 for 5 minutes. The fluid that remained on the brain was washed off with PBS. After the procedure, the skull was cleaned, the wound edge sutured and the mouse was put back into its home cage for recovery. Seven days later, brains were removed and processed for analysis as described below.

### Injections of retrobeads

Injections of retrobeads (Lumaflour) were made in S1 cortex in L1 at a depth of ∼150-200 microns from pia (200 nl, 20 nl/ min). The depth was chosen to be just under L1 in order to take advantage of backflow along the injection track. After the procedure, the skull was cleaned, the wound edge sutured and the mouse was put back into its home cage for recovery. Seven days later, brains were removed and processed for analysis as described below.

### Stereotaxic viral injections

For stereotaxic injections, an incision was made in the scalp, the skull was cleaned with PBS, and a small craniotomy was drilled above the injection location using a dental drill. The brain tissue was kept moist by applying sterile PBS at regular intervals. For injections, we tip-filled pipettes using negative pressure, and injected with pipettes of a tip size of approximately 4 µm. Viral injections -- 200 nl under constant positive pressure with a flow rate of 20 nl/min -- were made using the following stereotaxic coordinates for S1 cortex: AP -1.5 mm, lateral: 2.7 mm, ventral: 0.8 mm; for POm: AP: -1.5 mm, lateral: 1.5 mm, ventral: 3.0 mm.

To induce Cre specific expression of channelrhodopsin, an AAV1-Flex-ChR2-eYFP (1 x 10^13 vg/mL) was injected into S1 of Tlx3-Cre and Sim1-Cre mice. To induce rabies virus expression, we first injected 200 nl AAV-Syn-Flex-nGToG-WPRE3 (8.1 x 1011 GC/mL) into L5a and L5b and allowed two to three weeks for expression, followed by injection of 200 nl of RABVSADB19dG-mCherry (1.4 x 108 IU/mL). When combining rabies virus injection with fb application, we applied fb as on the day of rabies virus injection and waited seven days for expression. We analyzed every second section. In rabies virus injections, GFP positive neurons were distributed from 600 µm to 1100 µm across the injection site.

### Slice preparation for electrophysiology

We prepared coronal slices (300 µm) from Tlx3-Cre and Sim1-Cre mice. Immediately following the slice preparation, we used ice-cold solution for slicing and incubation of the brain slices for 5 min at 32 °C in (mM): 110 choline chloride, 2.5 KCl, 1.25 NaH2PO_4_, 26 NaHCO_3_, 11.6 sodium ascorbate, 3.1 sodium pyruvate, 7 MgCl_2_, 0.5 CaCl_2_ and 10 d-glucose, pH 7.4, saturation 95% O_2_/ 5% CO_2_, then the slices were incubated for a further 25 minutes at 32 °C in normal artificial cerebrospinal fluid (ACSF, in mM: 125 NaCl, 2.5 KCl, 1.25 NaH2PO_4_, 25 NaHCO_3_, 2 CaCl_2_, 1 MgCl_2_, and 25 dextrose, saturation 95% O_2_/ 5% CO_2_). ACSF was used for long-term incubation at room temperature and recordings in a submersion chamber at 32 °C.

### Definition of cell types for recording

We recorded from eYFP-negative neurons in L5 at the site of the viral injection (where expression was maximal). We classified neurons as L5b pyramidal cells if they had a single apical dendrite that extended to L1, their somata were about 800 µm from pia, and they had electrophysiological characteristics of pyramidal tract neurons, such as large sag current upon hyperpolarization and low input resistance. L5a pyramidal neurons had somata less than about 700 µm from pia, and a thin apical dendrite and small tuft in L1. Neurons with somata in L1 with non-spiny dendrites were classified as L1 interneurons.

### Recording and imaging

Whole-cell current clamp recordings with low resistance (4-7 MΩ) microelectrodes were made using a Dagan (Minneapolis, MN) BVC-700A amplifier. For current-clamp recordings the intracellular solution contained (in mM): 115 K-gluconate, 5 KCl, 10 HEPES, 4 ATP-Mg, 0.3 GTP-Tris, and 10 phosphocreatine, 0.1% biocytin. The pH was adjusted with KOH to 7.25-7.30 (285-295 mOsm). The liquid junction potential was not corrected. Slices were visualized on an Olympus (Tokyo, Japan) BX50WI microscope using a Photometrics CoolSnap ES (Tuscan, AZ) CCD camera and oblique optics. During recording or post-recording, neurons were filled, identified and categorized using Alexa Fluor 594 fluorescent dye (Invitrogen, Carlsbad, CA; 10 mM) and/or biocytin (0.2%).

### Channelrhodopsin based circuit mapping (sCRACM)

sCRAM was used to test the effect of Tlx3-Cre or Sim1-Cre input to L1 interneurons and to examine connections between L5 pyramidal neurons. We injected AAV1-Flex-ChR2-eYFP (1 x 10^13 vg/mL) into S1 of Tlx3-Cre and Sim1-Cre mice. Following three to five weeks of expression, we sliced the brains and recorded from neurons in L1 and L5 in the region of the densest ChR2-eYFP label. ChR2 photocurrents and EPSPs were distinguishable based on the response time during optical stimulation. Recordings of synaptic input were done in the presence of tetrodotoxin (TTX) and 4-aminopyradine (4-AP) to ensure that the connections being activated were monosynaptic.

### Glutamate uncaging

One photon glutamate uncaging was performed in brain slices taken from wildtype mice through a 60x/0.9 NA Olympus objective. Following baseline recording, slices were bathed in TTX (0.5 µM) and 4AP (1.2 mM), and MNI glutamate (4-Methoxy-7- nitroindolinyl-caged-L-glutamate, MNI-caged-L-glutamate, 0.2 µM, TROCIS), and neurons in L5a and L5b stimulated at three points along their dendritic axes. To evoke EPSPs stimulation, glutamate was uncaged for 30 ms duration with three optical pulses using 405 nm wavelength.

### Pharmacology

The recordings for sCRACM were done in the presence of TTX (0.5 µM) and 4-AP (1.2 mM) following baseline recordings. The recordings for glutamate uncaging were performed in addion with caged glutamate (4-methoxy-7-nitroindolinyl-caged-L-glutamate, MNI-caged-L-glutamate, 0.2 µM).

### Histology and Nissl stain

Mice were deeply anesthetized with isoflurane (Baxter Inc.), and transcardially perfused with 0.1% PBS for three minutes, and subsequently with 4% paraformaldehyde (PFA) in 0.1M phosphate buffer (PB, pH 7.2) for seven minutes. Brains were carefully removed from the skull and post-fixed in 4% PFA overnight at 4 °C. For sectioning with a vibratome, brains were embedded in 3% agarose, for sectioning with a microtome, brains were soaked in 30% sucrose overnight. All brains were sectioned at 70 µm. Every second section was mounted on slides and cover slipped. The remaining sections were stored in cryo-protect medium (30% Ethylene Glycol, 30% Glycerol in PBS) at –20 °C. We performed nissl stain with NeuroTrace^TM^ 500/525 Fluorescent Nissl Stain (Thermofisher, N21480) according to the protocol: Brain sections were washed in PBS three times, and then incubated in nissl solution (1:500) for 20 min. After the labelling step, brain sections were washed three times with PBS and mounted in glycerol (80%, 2.5% DAPCO in PBS).

### Confocal imaging

Images were taken on confocal laser scanning microscopes (Leica SP5, Leica SP8, Nikon A1Rsi+, Nikon Spinning Disk Confocal CSA-W1 SoRa), or on a widefield microscope (Nikon Widefield Ti2). Images were obtained with a 10x air, and 20x-, 40x-, or 63x oil immersion objectives (Leica: HCX PL APO 20x/0.7, HCX PL APO 63x/1.20 W motCORR CS; Nikon 20x Plan Apo, Air, 0.8 NA, 1.000 DIC N2 VC, Nikon 40x Plan Flour, Oil, 1.3 NA, 200WD, DIC N2 H; Nikon 40x Plan Flour, Sil, 1.25 NA, 300WD, SR HP DIC N1 λS OFN25; Nikon 10x Plan Fluo Air, 0.3 NA, 15.200 WD, Ph 1 DL; Nikon 10x Plan Fluo Air, 0.45 NA, 4.000 WD, DIC N1 λ OFN25). Fast blue was imaged with a 405 nm, GFP with a 488 nm, and tdTomato, mCherry or RFP with a 561 nm laser (bandwidth, Leica, 425/70, 515/25, 590/70; Nikon, 447/60, 450/50, 525/50, 595/50; Nikon widefield, 435/33, 519/26, 595/31).

### Definition of application and injection sites

For histological analysis, the application site for fb and the injection sites for retrobeads and rabies were defined by visual examination as the brain section with the densest labelling. Note that fb could spread from the application site at barrel cortex into lateral hindlimb / trunk S1 cortex (also see Cauller, 1991). For analysis we used every second brain section and stored alternate sections for additional examination.

### Cell counting and definition of cortical areas and cortical layers

Cell bodies were manually counted using the cell counter and analyze particles plugins from Fiji/ ImageJ. Labelling directly at the centre of the application site can be dense and individual neurons difficult to count, therefore we counted local labelled neurons at the centre of the application site and at 140 microns on each side of the centre of the application site where the fb label was less dense. For retrobeads, counts were made only at the injection site because label of tracer was less dense.

Cortical areas were defined according to the Allen brain reference atlas (https://mouse.brain-map.org/) and the Paxinos Atlas (Franklin and Paxinos 2008). The coordinates we used for each area were: M1 (AP 1.0 to 1.5, lateral 1.0 to 1.5), M2 (AP 2.0 to 2.5, lateral 1.0 to 1.5), S1 including trunk/ limb/ barrel field (AP -1.0 to -1.8, lateral 2.3 to 2.7), S2 (AP -0.5 to -1.8, lateral 2.7 to 4.2), visual cortex, including V1, V2L (AP -2.9 to -4.0, lateral 1.5 to 3.0), perirhinal cortex (AP -1.5 to -3.5, lateral 3.0 to 3.8).

Cortical layers in the brain areas were identified according to the Allen brain reference atlas and in bins of 100/ 200 microns for each layer. In S1 cortex, layers were identified according to these criteria, in microns from pia: L1 1-100 µm, L2/3 100-300 µm, L4 300-400 µm, L5a 400-500 µm, L5b 500-700 µm, L6a, 700-900 µm. Layers for M1 and M2 cortex were identified according to these criteria, in microns from pia: L1, 1-100 µm, L2/3, 100-300 µm, L5a, 300-400 µm, L5b, 400-600 µm, L6a, 600-800 µm. For S2 and visual cortices (including V1, V2L), we set the criteria for layers, in microns from pia: L1 1-100 µm, L2/3 100-250 µm, L4 250-350 µm, L5a 350-450 µm, L5b 450-600 µm, L6a 600-800 µm. In all these areas, L6b was also defined as the layer within 100 microns above the white matter. In our scheme L5a was 100 microns and L5b was 200 microns. Because laminar definitions are to some extent arbitrary (Oberlaender et al. 2011), to examine how our definition of layers affects our results, in a subset of brains, we also counted labelled neurons in a 30-micron strip in the middle of each 100-micron bin.

### Statistics

Statistical analyses were carried out with graphpad/ Prism. Data are shown as mean ± S.E.M. (standard error of the mean). p-values are indicated in the text: *p=0.05-0.01, **p=0.01-0.001, ***p=0.001-0.0001; ****p=<0.0001. For figure 3, a Fisher’s Exact test and Student t-test was used. For comparison between fb application and retrobead injection, the Kruskal-Wallis test was used. For all other figures, a one-way ANOVA, and Bonferroni post hoc test was used.

## Results

### Local input to L1 of S1 cortex

To assess input to L1 of primary somatosensory cortex, the retrograde tracer fast blue was applied on the cortical surface (L. J. Cauller et al. 1998; Clancy and Cauller 1999; Keizer et al. 1983). Seven days after application of the tracer, labelled neurons were evident at the application site and in sites distributed throughout the brain (**Figure 1A**). The laminar pattern of the local uptake was specific and in agreement with the earlier qualitative observations in rat somatosensory cortex (L. J. Cauller et al. 1998; Clancy and Cauller 1999). Because label under the application site can be dense and individually labelled neurons difficult to distinguish from background, we assessed counts both at the centre of the application site and 140 microns from the centre of the application site. At 140 microns from the center of the application site, we found the highest uptake of fb in L2/3 (40% of all fb labelled neurons in n=6 sections from n=6 mice) and in L5 (L5a: 18%, L5b: 11%). L4 and L6a had a smaller percentage of labelled neurons (L4, 5%, L6a, 1%), and L6b had 13% of fb labelled neurons. The proportion of total fb labelled neurons in L1 was 13%. When we compared the counts 140 microns from the centre of the application site to counts directly under the centre, we found that the laminar pattern of local labelled neurons was similar (n=6 mice, 3096±459 fb labelled neurons per S1 cortex, one-way ANOVA ****p<0.0001, **Figure 1B**, **Table 1A, Supplementary Figure 1A-C, Supplementary Figure 2A)**.

**Figure 1.**
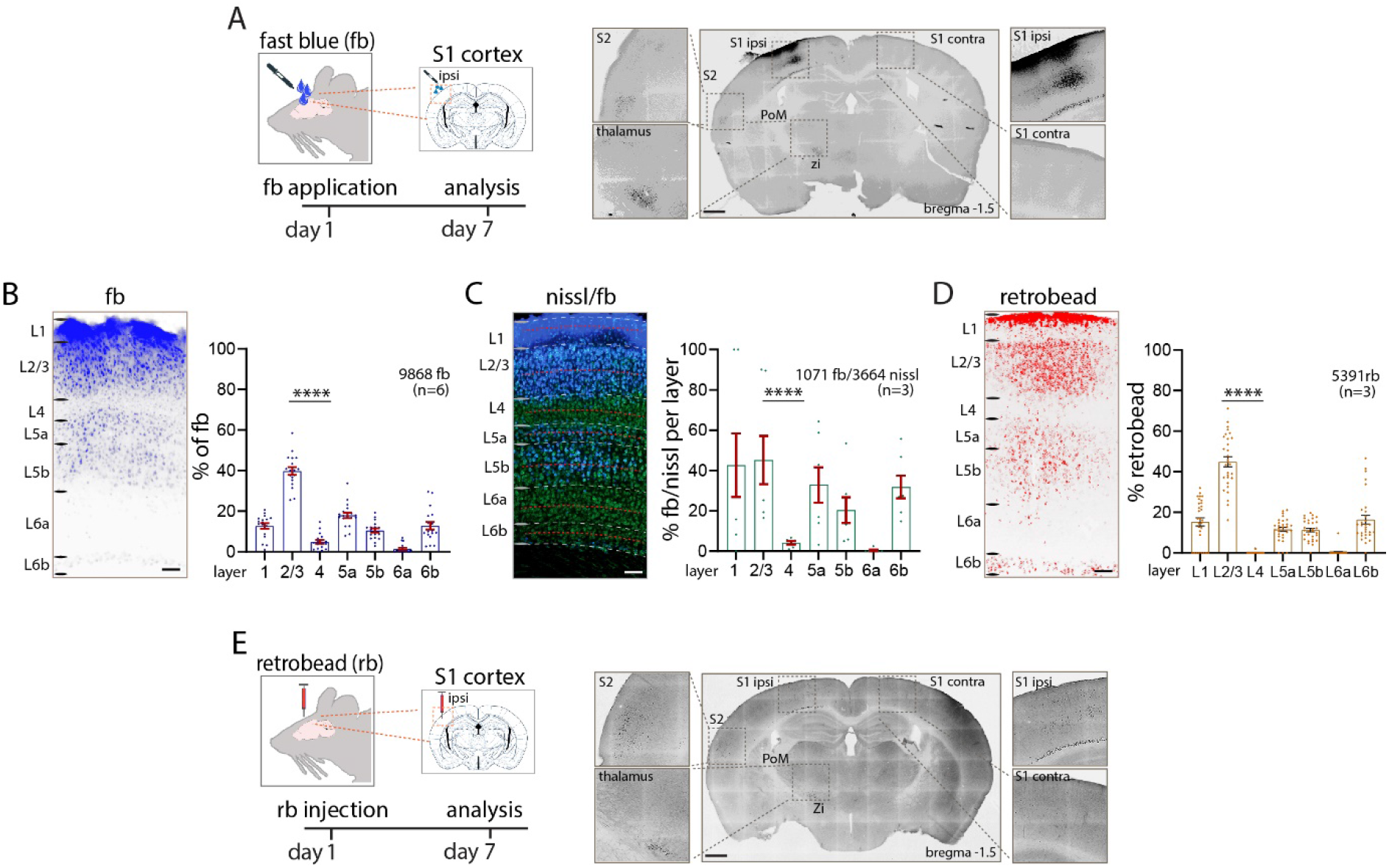
Local Input to L1. **(A)** Schematic of experiment. Retrograde tracer fb was applied on L1 of S1 cortex and fb labelled neurons in S1 cortex were counted seven days after incubation. **(B)** Uptake of fb was evident across all cortical layers, with prominent uptake in L2/3 and L5. The tabulation of fb labelled neurons in different lamina under the application site is shown in the bar graph. **(C)** Percentage of L1 projecting fb labelled neurons in each layer. Nissl stained (green) and double labelled (fb and nissl) neurons were counted to obtain a percentage of labelled neurons in each lamina (30-micron bin, red dotted line). L2/3 and L5 had the highest percentage of double labelled neurons. **(D, E)** Injection of retrobeads in L1 shows a similar pattern of local and long-range input to L1 as with fb application. Total number of neurons counts are shown in each panel. Statistical analysis with one-way ANOVA, Bonferroni post-hoc test, ****p<0.0001. Total number of neurons counted and mice used (in brackets) are shown in each panel, analysis details in **Table 1A**. Scale bars in **A, E** 500 µm, in **B, C, D** 100 µm.

The percentage of neurons labelled with fb in each layer was assessed by combining Nissl staining with fb application. The number of double labelled (nissl and fb) neurons in a 30-micron bin in the middle of each defined layer were counted and the laminar pattern assessed. These analyses showed that 45% of all neurons in L2/3, 33% of all neurons in L5a and 20% of all neurons in L5b were double labelled. Few neurons in L4 (4%) and L6a (0.4%), and 32% of L6b neurons were double labelled. Finally, 43% of neurons in L1 were double labelled (n=6 mice, n=3 nissl + fb stained sections, one-way ANOVA **p<0.01; **Figure 1C**, **Table 1A**). The laminar pattern of fb with relation to nissl stain was similar when we used 100 micron bins to define each layer (**Supplementary Figure 1D**).

Next, we assessed input to L1 with retrobeads injected into cortex. When retrobeads were injected in L1, we obtained a similar laminar pattern as with application of fb on the surface of cortex (n=3 mice, 2490±374 retrobead labelled neurons per S1 cortex, **Figure 1D, 1E, Supplementary Figure 2B, Table 1A**). Locally, 45% of all retrobead labelled neurons were in L2/3, followed by 12% of L5 neurons (L5a: 12%, L5b: 11%). We found little uptake of retrobeads in L4 (0.1%), no label in L6a (0.3%), 16% of neurons labelled in L6b and 15% in L1. We found no significant differences in the laminar distribution of fb labeled versus retrobead labeled neurons at the application / injection site (**Table 1A**). From these experiments, we conclude that input to L1 has a specific local laminar pattern that can be assessed by fb or retrobeads, with the majority of local projections arising from L2/3, L5 and L6b.

### Long-range input to L1

To estimate the laminar distribution of long-range input to L1 of S1 cortex, we counted and categorized fb labelled neurons in a variety of cortical areas (**Figure 2, Supplementary Figure 3, Table 1B**).

**Figure 2.**
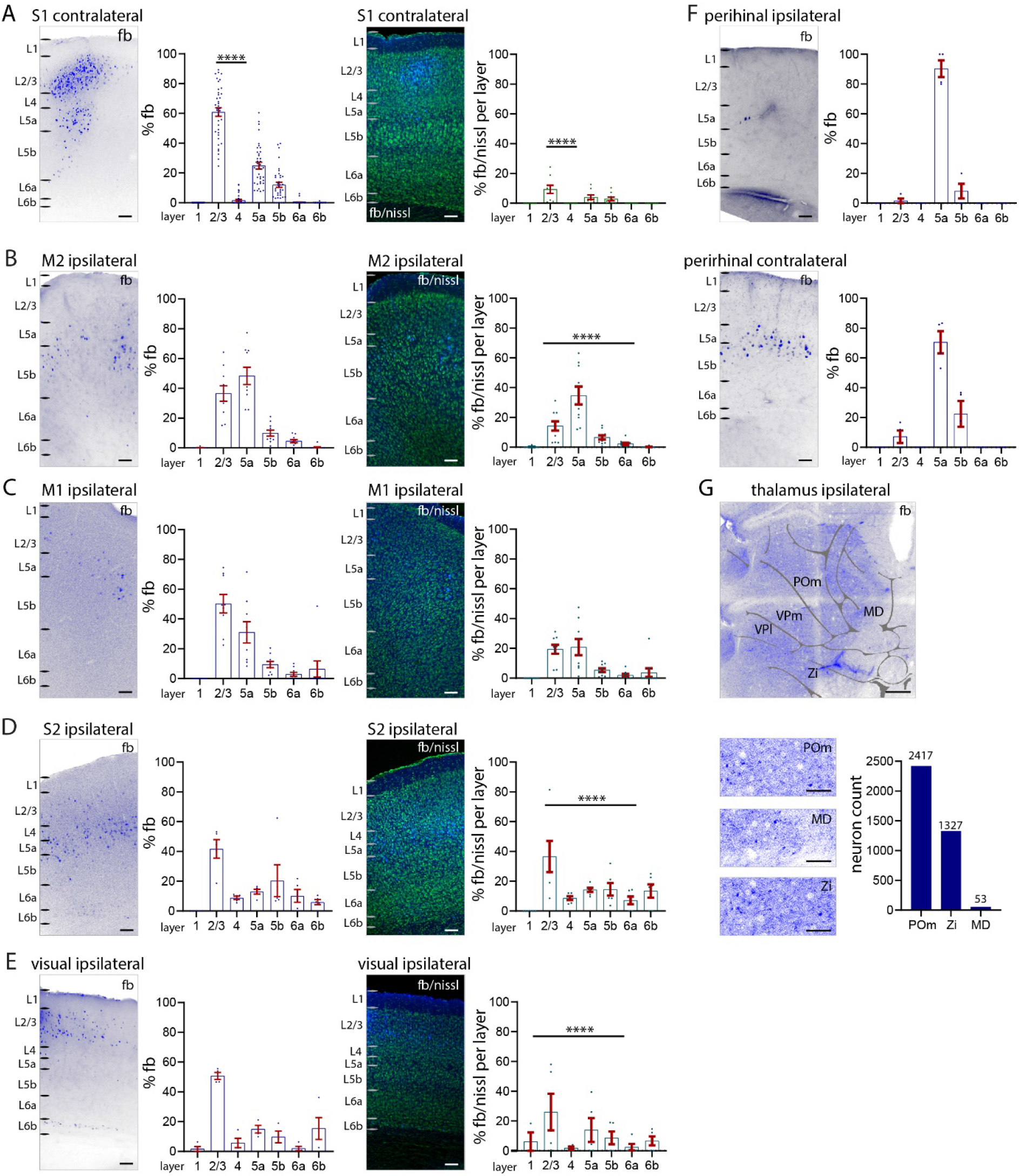
Long-range input to S1 L1 from other cortical areas. **(A-E)** Percentage of total fb labelled neurons (*left*), and percentage of labelled neurons in relation to nissl in each layer (*right*). (**A**) S1 contralateral cortex (**B**) M2 cortex, (**C**) M1 cortex, (**D**) S2 cortex, (**E**) visual (V1, V2L) cortices, (**F**) perirhinal cortices ipsilateral and contralateral. Counts of fb labelled neurons ipsilateral to application site (left) and contralateral (right). Quantification of the laminar pattern of long-range input to L1 shows the highest percentage of fb labelled neurons in cortical L2/3 and L5a. (**G**) Fast blue label (total numbers) in POm thalamus, zona incerta and midline thalamus shows that these three subcortical nuclei have projections to L1. Abbreviations in **G**: MD, midline; POm, posteromedial complex of thalamus; VPl, ventrolateral nucleus of thalamus; VPm, ventromedial nucleus of thalamus; Zi, zona incerta. Each dot in the graphs represents one brain section. Data are from four brains. Statistical analysis with one-way ANOVA, Bonferroni post-hoc test, ****p<0.0001. Analysis details in **Table 1B**. Scale bars in **A-F** 100 µm, in **G** 500 µm, in zoom-ins 50 µm.

Fast blue labelled neurons were evident in contralateral S1. The highest percentage of fb labelled neurons were found in L2/3 (61% of all labelled neurons), fewer were found in L5 (L5a: 25%, L5b: 12%). Very few neurons in L4 (1.7%) and L6a (0.3%) were labelled. No neurons in L1 and L6b were labelled. The total number of fb labelled neurons in contralateral S1 (366±55 neurons per brain) were 10% of the total number of fb labelled (3096±459 neurons per brain, see **Figure 1**) neurons in the ipsilateral S1 (one-way ANOVA ****p<0.0001). Overall, our analysis of fb and nissl stained neurons shows that 9% of all L2/3 neurons project to the contralateral L1 S1. Four percent of L5a and 3% of L5b neurons were retrogradely labelled and projected to L1 of contralateral S1. Few neurons in L1, L4, L6a, or L6b were fb labelled or projected to L1 of contralateral S1 (**Figure 2A)**.

In higher order motor cortices (M2), most fb labelled neurons were in L2/3 and L5a, with a significantly larger number of fb labelled neurons in L5a (35%) than in L2/3 (14%) (one-way ANOVA ****p<0.0001; **Figure 2B**). In M1, most fb labelled neurons were distributed roughly equally in L2/3 (19%) and L5a (21%) (**Figure 2C**). In S2 cortex, most fb labelled neurons were located in L2/3 (42%), there were a smaller number of fb labelled neurons in other layers (**Figure 2D**). In visual cortices (including V1 and V2L), most fb labelled neurons were located in L2/3 (42%), followed by L5a (23%) (**Figure 2E**). In perirhinal cortices, neurons in L5a from both hemispheres project to S1 L1 (Doron et al. 2020), with the bulk of the perirhinal projection to L1 of S1 from the contralateral hemisphere. The perirhinal input arose from neurons in L5a (**Figure 2F**), which accounted for 2% of neurons in L5a in ipsilateral and 22% in contralateral perirhinal cortex.

Fast blue labelled neurons were also found in subcortical areas, POm, zona incerta, midline thalamic nuclei, basal forebrain and brainstem. Subcortically we only counted the number of fb labelled neurons in the thalamus and zona incerta **(Figure 2G)**. We made no attempt to determine the percentage of labelled neurons in each nucleus (i.e. the ratio of fb labelled neurons to nissl stained cells).

When fb was injected into cortex -- label was distributed from L1 to L6 at the injection site -- the laminar pattern of long-range input to S1 was statistically, significantly different from that observed with fb application on the cortical surface (**Tables 1A, 1C**). The cortical areas containing retrogradely labelled neurons were the same when fb was injected into cortex or applied on the cortical surface. But in contralateral S1, injections of fb revealed more labelled neurons in L6a that were not evident when fb was applied on the cortical surface. Note that whether fb was applied on cortex or injected in cortex, most of the retrogradely labelled neurons in contralateral S1 were in L2/3. In S2, the largest number of fb labelled neurons after injection of fb were found in L2/3 and L6a (one-way ANOVA, *p<0.05). In motor cortices, fb neurons were distributed across all layers (**Supplementary Figure 4A-F**). These results suggest that long-range projections to L1 consist of a specific set of neurons located in L2/3 and L5. In contrast, long-range projections to deeper parts of S1 engage more neurons in L6a in both contralateral S1 and ipsilateral S2.

The injections of fb also served as a control. Because VPm thalamic axons target L4 and can even target pyramidal cell dendrites in L2/3 of cortex, label in VPm or VPl thalamus is only expected if fb applied on cortical surface seeps into L2/3 or L4, or after injection of fb into cortex (Guest et al. 2021; Killackey and Ebner 1973; Lu and Lin 1993; Sermet et al. 2019). In experiments with fb application on cortical surface, we therefore excluded brains that showed fb label in VPm and VPl because label at these sites would indicate that fb had seeped into L2/3 or L4 at the application site (**Supplementary Figure 1E-1F**). Application of fb in Scnn1a-Cre mice (Madisen et al. 2010) confirmed that fb was not labeling cells in L4 (**Supplementary Figure 1G**). These results confirm that local laminar distribution of fb label was significantly different when fb was applied on L1 compared to when fb was injected into cortex (**Tables 1A, 1C**).

### Functional effects of L5 monosynaptic connections in L1

The local pattern of fb distribution in S1 cortex suggest that a substantial input to L1 arises from local L2/3 and L5 neurons. The synaptic connectivity between L2/3 neurons and neurons in L1 has been examined previously (Wozny and Williams 2011), so here we focused on the effect of L5 inputs to L1. We examined the synaptic connectivity between L5 pyramidal neurons and L1 interneurons as well as between L5 pyramidal neurons themselves (via connections within L1). We patched L1 interneurons or L5 pyramidal neurons in Tlx3-Cre or Sim1-Cre mice expressing ChR2. To examine 1) input to L1 interneurons, we photo-stimulated over L1 while recording from L1 interneurons using normal ASCF solution without TTX and 4-AP. To examine 2) connectivity between L5 pyramidal neurons, in the same experiment, we then photo-stimulated at different points along the somato-dendritic axis of L5 pyramidal neurons in the presence of TTX and 4-AP. This approach blocks di-synaptic transmission (Petreanu et al. 2007). We expected the IT input to generate a larger PSP than the PT input, in part because more IT neurons project to L1. The PSP parameters -- amplitude, latency, and short-term plasticity (at 10 Hz) -- evoked in interneurons and in L5 pyramidal neurons – were measured but due to the use of TTX and 4-AP, which can alter normal release characteristics, these were not characterized in depth.

#### Effect of L5 input on L1 interneurons

We recorded from interneurons in L1 in Tlx3-Cre or Sim1-Cre slices while photo-stimulating in L1. This experiment was performed without TTX and 4-AP (**Figure 3A**). Photo-stimulation of the axons in L1 showed that IT neurons generate a powerful PSP in L1 interneurons and that this input was significantly stronger than the PT input to the interneurons (**Figure 3B, 3C**; n=5 neurons, Tlx3 to L1 interneuron amplitude: 25.5±4.9 mV; Sim1 to L1 interneuron amplitude: 0.9±0.2 mV, ****p<0.0001, Student t-test). Also, note that five out of five interneurons received input from IT while only two out of five interneurons received input from PT. The input from IT to L1 interneurons had depressing synaptic dynamics at 10 Hz stimulation (**Figure 3D**).

**Figure 3.**
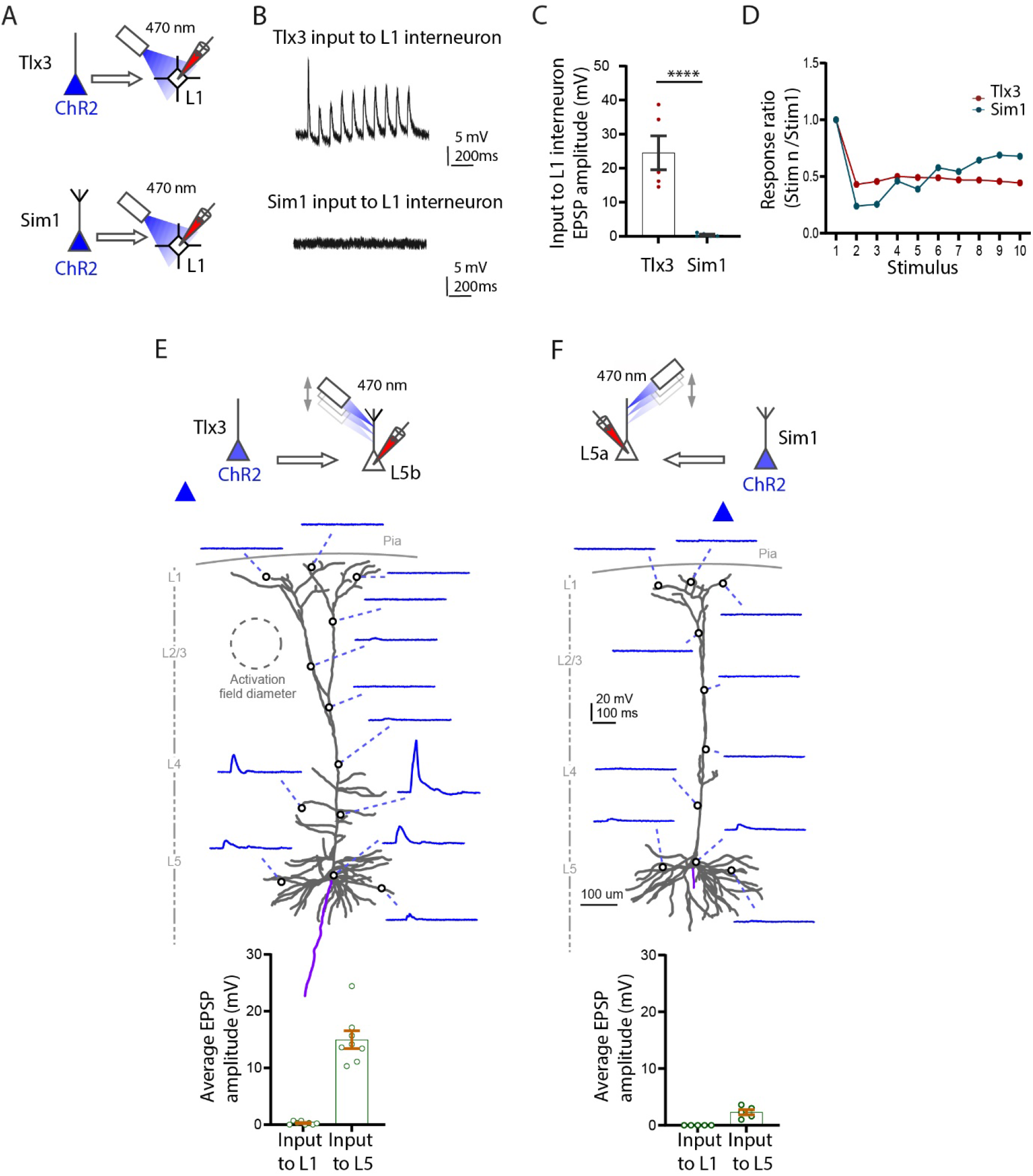
Functional synaptic input locally to L1 from L5 Tlx3 and Sim1 neurons in S1 cortex. (**A**) Experimental setup to test the input from L5 Tlx3-Cre and L5 Sim1-Cre neurons onto L1 interneurons. (**B**) Example recordings of input to L1 interneurons from Tlx3 (*top*) and Sim1 (*bottom*) neurons. (**C**) Group data of evoked EPSPs in L1 interneurons (Tlx3, n=5; Sim1, n=5, three mice each genotype). (**D**) Short-term dynamics of evoked EPSPs in L1 interneurons (10 Hz; Tlx3, n=5; Sim1, n=5, three mice each genotype). (**E, F**) *Top*, experimental setup to test the input from L5 IT (Tlx3) and L5 PT (Sim1) neurons onto subcellular spots along L5b PT (n=8, three mice) and L5a IT (n=5, three mice) neurons, respectively. *Middle*, example neurons and their responses to light spot illumination in the indicated locations on each neuron (**E**, L5b neuron, input from Tlx3; **F**, L5a neuron, input from Sim1). *Bottom*, group data comparing the input to L1 (apical tufts) and L5 (somatic region). **A-D**, experiments performed in normal ACSF solution; **E, F**, experiments in the presence of TTX and 4-AP.

#### IT-PT interactions with other L5 pyramidal neurons

Next, we examined whether IT neurons could drive non-Tlx3 neurons in L5b, and whether PT neurons could drive non-Sim1 neurons in L5a (**Figure 3E, 3F** *schematic top*). In order to restrict the transmitter release to the photo-stimulated area, and to prevent di-synaptic activation, here we applied TTX and 4-AP to the bath solution. The entire somato-dendritic axis of the filled neurons was systematically photo-stimulated by moving the optogenetic light in 50 µm steps, while simultaneously measuring the amplitude of the evoked EPSP. Activation of the Tlx3-Cre axons did not evoke any EPSPs in L5b tufts in L1. Similarly, activation of Sim1-Cre axons did not evoke EPSPs in non-Sim-L5a tufts in L1 (**Figure 3E**, *bottom graph*). The Tlx3 neurons did, however, evoke a significant depolarization (while photo-stimulating) at the oblique dendrites and to a lesser extent at the basal dendrites and soma in L5b (14.97±1.56 mV). The Sim1-Cre axons on the other hand, had a negligible effect on non-Sim L5a neurons across the entire somato-dendritic axis (2.30±0.46 mV) (**Figure 3F**, *bottom graph*) (Ramaswamy et al. 2012).

In summary we show that the L5 Tlx3 IT neurons strongly drive interneurons in L1 but have a negligible effect on the apical tuft dendrites in L1 of non-Tlx3 L5b neurons. Layer 5 Sim1 PT neurons do not drive interneurons in L1 or the tuft dendrites of L5 non-Sim1 neurons in L1. This suggests that two major classes of L5 pyramidal neurons do not interact with each other directly within L1.

#### Glutamate uncaging

To confirm that the input to the apical tuft dendrite could evoke a synaptic response in our experimental conditions, we performed glutamate uncaging along the dendritic axis while recording from neurons in L5a and L5b (**Supplementary Figure 5A, B**). Our results indicate that the effect of localized input at the tuft dendrite can be measured at the soma. Glutamate uncaging evokes measurable EPSPs in both cell types at the soma, when stimulating a local spot in the apical tuft dendrite for 30 ms in the presence of TTX/ 4-AP. Our work and previous work (Hooks et al. 2011; Passlick and Ellis-Davies 2019) indicates that our methods using sCRACM are potentially sensitive enough to detect inputs to the apical tuft dendrites of L5 cells.

### Cell type specific projections to S1 L1

To assess the cell type specificity of input to L1, we applied fb on the surface of L1 in somatosensory cortex in Ai9-reporter mice. We used Tlx3-Cre mice for L5 intratelencephalic (IT) neurons, and Sim1-Cre mice for L5 pyramidal tract (PT) neurons (Gerfen et al. 2013; Gong et al. 2003). For subclasses of L6b projection neurons we used Ctgf-2A-dgCre and Drd1a-Cre mice (Heuer et al. 2003; Hoerder-Suabedissen et al. 2018; Zolnik et al. 2020). To examine the inhibitory component of input to L1, we used Vip-Cre for vasoactive intestinal peptide, Sst-Cre for somatostatin neurons (Taniguchi et al. 2011), and PV-Cre for parvalbumin neurons (Hippenmeyer et al. 2007).

Layer 5 and L6b neurons in S1 cortex. We first assessed the percentages of layer specific cell types for each Ai9-reporter line (**Supplementary Figure 6A, 6B, Table 2A**). Tlx3-Cre neurons were exclusively found in L5, primarily in L5a. They constitute 42% of nissl stained neurons in L5, with 76% in L5a, and the remaining 24% in L5b. By comparison, Sim1-Cre neurons were distributed from L4 to L6, with 54% in L5b, and 28% in L5a and the rest (18%) in other layers. Overall, Sim1-Cre neurons constituted 18% of all neurons in L5. Note that although tdTom positive neurons are found across L5 in both Cre-lines Tlx3-Cre and Sim1-Cre, the majority of tdTom positive neurons for Tlx3 were in L5a, and the majority tdTom positive neurons in Sim1 were in L5b. Drd1 positive neurons were found in L6a and L6b, with the majority in L6b.

All Ctgf neurons were located in L6b, but Ctgf neurons constituted only 15% of all neurons in L6b. By comparison, Drd1a positive neurons were distributed in L6a and L6b, with 76% of all Drd1a neurons in L6b. Drd1a positive neurons were 43% of all neurons L6b (**Supplementary Figure 6C, 6D).**

In S1 cortex, in our Vip-, Sst-and Pv-Cre lines 13% of neurons were Sst positive (3220 tdTom out of 23889 nissl), 5% were Vip positive (460 tdTom out of 9780 nissl), and 14% were Pv positive (3936 tdTom out of 28899 nissl). These neurons were distributed in all cortical layers (**Supplementary Figure 6E-6G, Table 2B).**

High magnification images of Sim1-Cre-Ai9 show fb/ tdTom processes and neurons in L1 and L5b. These images show that adjacent tdTom positive neurons that have dendrites extending toward L1, do not necessarily take up fb, i.e. their somas are not uniformly fb positive (**Supplementary Figure 6H**). These images argue for a specific retrograde uptake of fb. Local projections from layers 5, 6b, and from inhibitory neurons to L1. To examine the IT and PT connectivity to L1, we applied fb on L1 in Tlx3-Cre-Ai9 and in Sim1-Cre-Ai9 mice and counted double labelled neurons in those lines (**Figure 4A, 4B, Supplementary Figure 6I, Tables 2C, 2E**). In ipsilateral S1, 29% of all IT neurons in L5 were fb labelled, with the majority located in L5a. The PT neurons projecting to L1 showed a different profile, with 20% of all PT neurons double labelled for fb ipsilaterally, with the majority of L1 projecting neurons located in L5b. In L6b, a larger proportion, 52%, of Ctgf neurons projected to L1, while only 13% of Drd1a neurons projected to L1 (**Figure 4C, 4D, Table 2C**). Fast blue labelled Sst neurons were most abundant in L2/3 and L5a, fb labelled Vip neurons were most abundant in L2/3, and fb labelled Pv neurons were most abundant in L5b (**Figure 4E-4G**, **Table 2D**).

**Figure 4.**
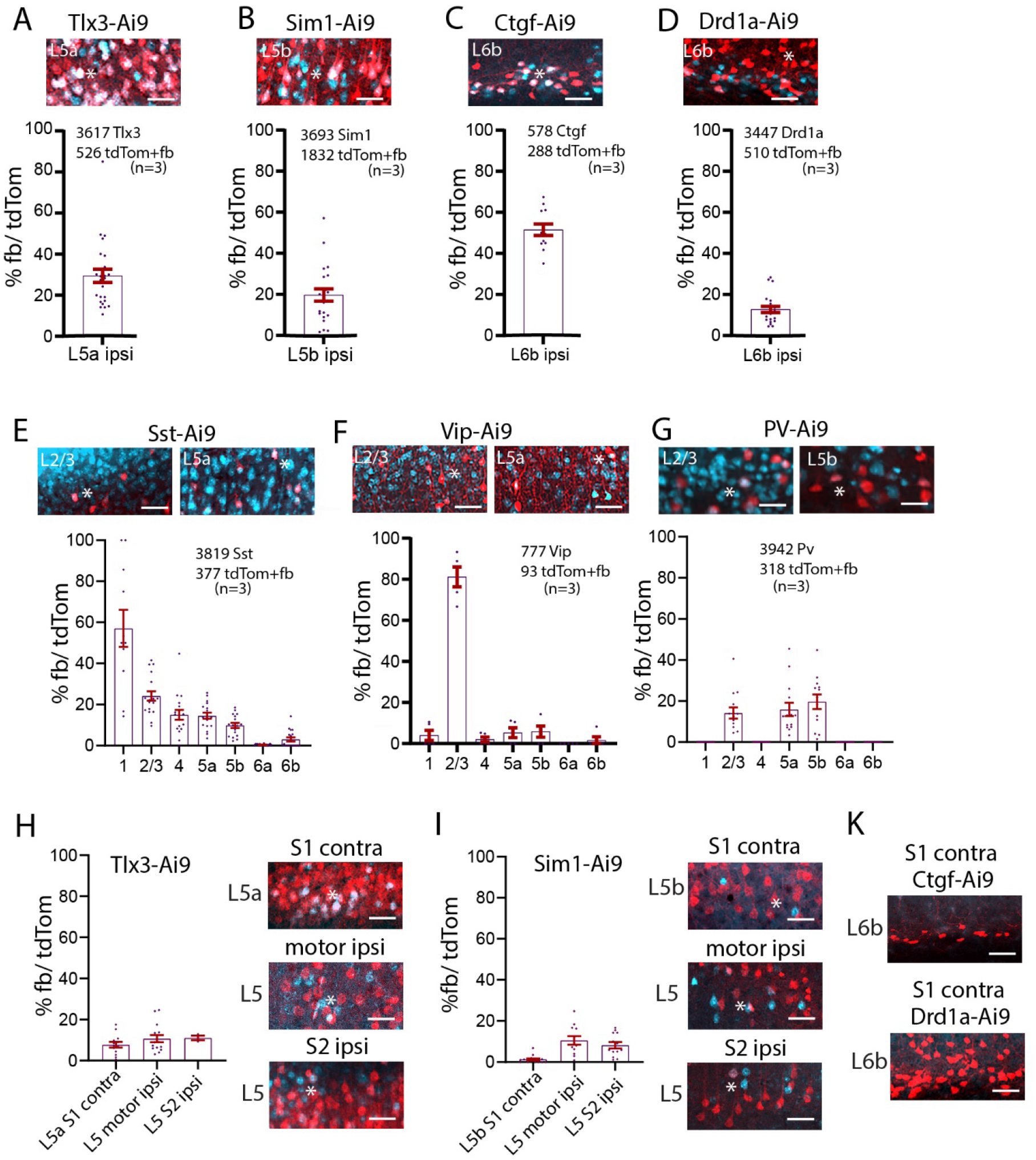
Local and long-range input from classes of L5 and L6b, and inhibitory cells to L1. **(A-G)** Example images of fb uptake in Tlx3-Cre, Sim1-Cre, Ctgf-Cre, Drd1a-Cre, SST-Cre, VIP-Cre, and Pv-Cre brains. Ipsilaterally, 29% of Tlx3 S1 project to L1. 20% of Sim1 neurons project to L1. 52% of L6b Ctgf positive neurons project to L1. Only 13% of L6b Drd1a neurons project to L1. SST and VIP neurons in all layers take up fb on the ipsilateral side. More than 80% of L2/3 Vip neurons take up fb. Pv neurons take up fb in layers 2/3 and 5. (**H)** Example images of long-range fb uptake in Tlx3-Cre brain. Contralaterally, 8% of Tlx3 neurons in S1 project to L1. Tlx3 neurons in motor and S2 cortices also project to L1. Quantification shown in graphs. **(I)** Example images of long-range fb uptake in Sim1-Cre mice. Contralaterally, few Sim1 neurons in S1 project to L1. Sim1 neurons in motor cortices and S2 cortices also project to L1. Quantification shown in graphs. **(K)** No Ctgf or Drd1 neurons contralaterally of S1 project to L1. Each dot in the graphs represents one brain section. Total number of neurons counted in each mouse line and mice used (in brackets) are shown in each panel, analysis details in **Tables 2C, 2D**. Fast blue pseudo colored in cyan. Scale bars 50 µm.

### Cell type specific long-range projections to S1 L1

IT and PT neurons in contralateral S1, ipsilateral motor cortices and S2 cortex were labelled with fb (**Figure 4H, 4I, Table 2C**). For contralateral S1, 8% of IT neurons, and only 1% of PT neurons were labelled. The percentage of double labelled neurons in motor cortices was similar for IT and PT neurons (∼10%). In S2, 11% of IT neurons, and 8% of PT neurons were double labelled. Thus, IT neurons project contralaterally to L1, and slightly more IT than PT neurons in motor and S2 cortices have long-range projections to L1. Overall, 1.5% of the IT neurons in perirhinal cortex project to contralateral S1 L1. No Ctgf or Drd1a positive fb double labelled neurons were found in the contralateral hemisphere (**Figure 4K**). In Sst projecting inhibitory neurons there were also a small number of long-range in contralateral S1, in L2/3 and L5a that targeted L1 (0.3%, 15 out of 6173 neurons, 3 brains).

### Local input to L1 and L5 neurons assessed with fb and rabies virus

The experiments so far revealed that subtypes of L5 neurons project in a specific manner to L1. These observations raise a question: Are neurons that project to L1 presynaptic to the L5 pyramidal neurons? To address this question, we combined fb application in L1 with rabies-based retrograde monosynaptic tracing for the IT and PT neurons. Monosynaptic rabies tracing labels neurons connected by one synapse to the source (starter) neurons (Kim et al. 2016; Luo et al. 2008; Wickersham et al. 2007).

#### Distribution of input along the somato-dendritic axis

To first examine whether the rabies virus targets synapses along the entire soma-dendritic arbor of the starter neuron -- i.e. to examine whether boutons of presynaptic neurons could be distributed in L1 to L6 -- we used a modified rabies virus linked to synaptophysin (Wickersham et al. 2013). With this approach, presynaptic cell bodies and their axons / boutons were labelled with synaptophysin linked to RFP (presynaptic) and the starter population was additionally labelled with cystolic GFP (postsynaptic) and nuclear cerulean fluorescent protein (CFP) (**Supplementary Figure 7A**). We examined the distribution of boutons and presynaptic (RFP) label in a circuit where it was known that a class of presynaptic neurons target specific dendritic loci. For this assay, we used the Gpr26-Cre line, which expresses Cre in a class of CA1 pyramidal neurons (**Supplementary Figure 7B**). Entorhinal neurons target the apical tuft dendrites of CA1 pyramidal neurons (Masurkar et al. 2017), so finding presynaptic, RFP labelled neurons in entorhinal cortex would indicate that the rabies strategy is effective even for inputs that target the tuft dendrites. When we examined entorhinal cortex, we found presynaptic, RFP labelled neurons, suggesting that rabies retrograde approach can work for synapses at the tuft dendrites (**Supplementary Figure 7C**).

Next, we used the same approach in S1 cortex of Sim1-Cre and Tlx3-Cre mice (**Supplementary Figure 7D-F**). Starter cells expressing CFP were found in L5, and GFP expressing cells and RFP positive boutons were detected in all layers, including L1, across the cortical column (Tlx3-Cre, n=369 synapses, Sim1-Cre, n=158 synapses, bouton counts from 10 boxes for each layer with a size of 20 µm x 10 µm). Presynaptic neurons were also detected in motor cortex, S2 and thalamus (**Supplementary Figure 7F**). These experiments suggest that the rabies approach targets the entire somato-dendritic compartment -- including synapses on apical tuft dendrites in L1 -- of the starter pyramidal neurons.

#### L1 projecting neurons, presynaptic to PT and IT neurons

With these results in hand, we performed injections in Tlx3-Cre and Sim1-Cre mice with the standard AAV-rabies virus approach in combination with fb application to see whether neurons that were presynaptic to L5 neurons also project to L1 (**Figure 5A, Supplementary Figure 8A, B**). In Tlx3-Cre mice, fb application in L1 labelled 9596 neurons, mCherry was found in 6138 cortical neurons presynaptic to the Tlx3 neurons. There were 449 neurons that were double labelled, i.e. were presynaptic to Tlx3 neurons and projected to L1. In Sim1-Cre mice, there were 13131 fb labelled, 6495 presynaptic neurons, and 823 double labelled neurons, i.e. those that were presynaptic to Sim1-Cre neurons and projected to S1 L1. The percentage of fb+/presynaptic was7.5% in Tlx3 and 9.0% in Sim1, the percentage of fb+/starter was 0.9% in Tlx3 and 5.2% in Sim1 (**Figure 5B**, **Table 3A, 3B**).

**Figure 5.**
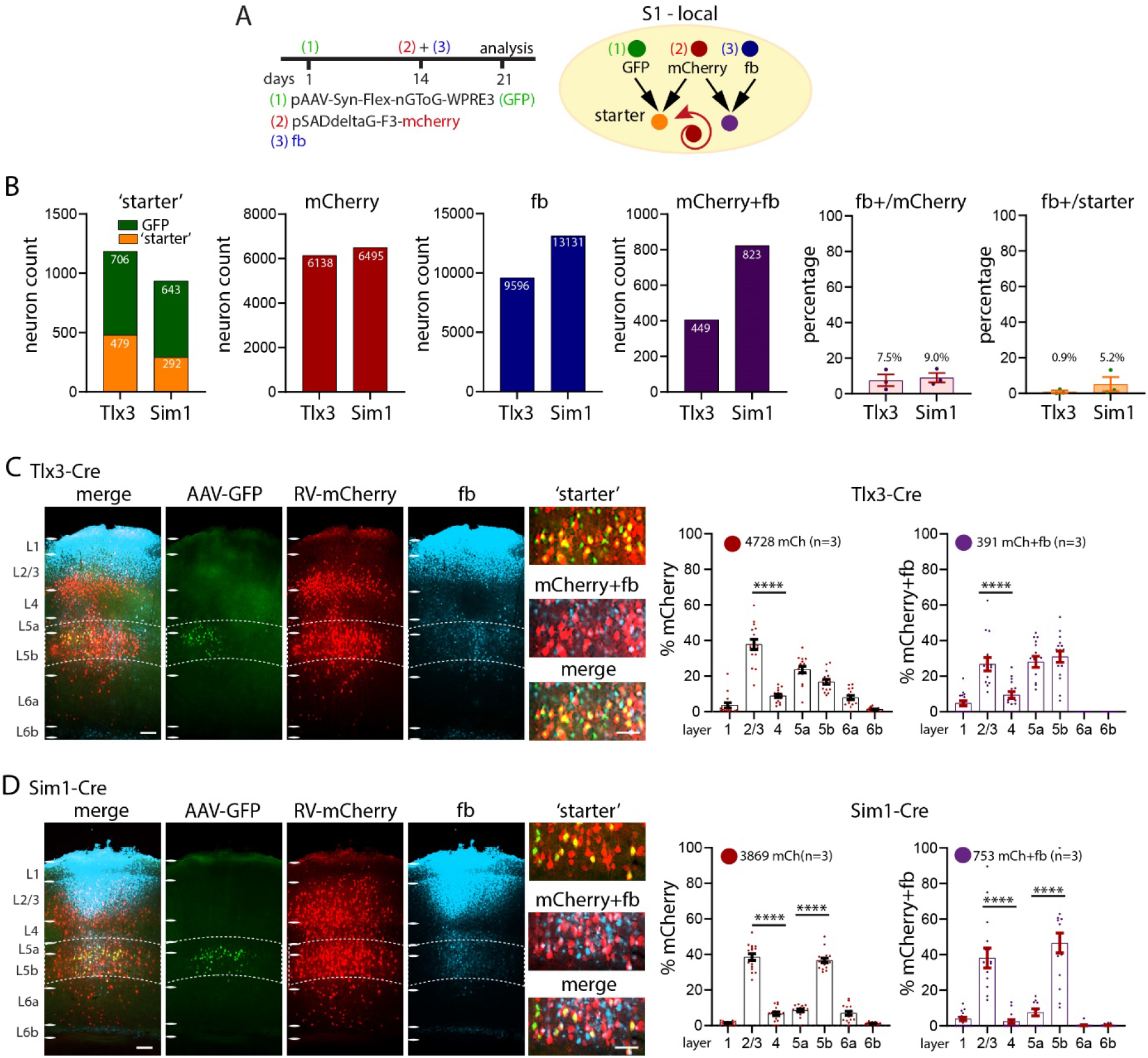
Local synaptic input to S1 L1 in Tlx3 and Sim1 neurons in combination with fb. **(A)** Schematic of injection scheme. Fourteen days after the first injection with AAV-Syn-flex-nGToG-EGFP (step 1), fb was applied on L1 in parallel to the injection of rabies virus (steps 2 and 3). Both, rabies virus and fb were allowed to express for seven days. (**B**) Number of starter, mCherry (presynaptic), fb labelled and double labelled neurons (mCherry+fb), and percentages of double labelled/ starter, and double labelled/ presynaptic in single mice. In Tlx3-Cre mice, the presynaptic neurons derived from 479 starter neurons (out of 706 GFP expressing neurons), in Sim1-Cre mice the presynaptic neurons derived from 292 starter neurons (out of 643 GFP expressing neurons). (**C, D**) Example images of virus expression and fb uptake in S1 cortex of (**C**) Tlx3-Cre and (**D**) Sim1-Cre brains. Insets showing starter neurons, presynaptic neurons and fb double labelled neurons. Percentages of presynaptic neurons in S1 cortex and double labelled fb and rabies, neurons in Tlx3-Cre and Sim1-Cre brains in graphs. Each dot in the graphs represents one brain section. Total number of neurons counted are shown in each panel, data from three mice each genotype, data shown as mean ± S.E.M. Statistical analysis with one-way ANOVA, Bonferroni post-hoc test, ****p>0.0001. Analysis details in **Tables 3A, 3B.** Scale bars in **C, D** 100 µm, in zoom-ins 50 µm.

#### Local intra-cortical pattern

Total presynaptic input was 6138 neurons in Tlx3 mice and 6495 neurons in Sim1 mice. Example images for the injection site in Tlx3-Cre and Sim1-Cre brains show starter cells in yellow and mCherry+fb double labelled cells in cyan. Locally, in Tlx3-Cre brains, the presynaptic labelled neurons were mainly seen in L2/3 and L5 (one-way ANOVA, ***p>0.001). Neurons that were double labelled i.e, were presynaptic to Tlx3 neurons and labelled with fb, were most common in L2/3 and L5a and L5b (**Figure 5C**). In Sim1-Cre brains, most of the local presynaptic input was from L2/3 and L5b (**Figure 5D**) and double labelled neurons were detected mainly in L2/3 and L5b.

#### A control for fb, when combined with rabies approach

We obtained similar results for fb application on L1 in the transgenic lines as we do with fb application in wild type mice (**Supplementary Figure 8C-F**). To confirm whether fb application on L1 interferes with rabies, we applied rabies virus to Tlx3-Cre and Sim1-Cre brains without fb application (**Supplementary Figure 9A-D, Table 3C**). The pattern of fb distribution was similar in both approaches. In Tlx3-Cre brains, presynaptic neurons were mainly found in L2/3 and L5a, in Sim1-Cre brains presynaptic neurons were mainly found in L2/3 and L5b.

### Long-range input to L1 and L5 neurons

Earlier work has shown that in addition to targeting L1, long-range input from cortical areas like M1 targets infragranular layers (Geng et al. 2022; Kinnischtzke et al. 2016; Yamawaki et al. 2021; Zagha et al. 2013). This raises the question whether neurons that project to L1 from other brain areas are presynaptic to the L5 pyramidal neurons? Long-range presynaptic input to Tlx3-Cre and Sim1-Cre neurons in S1 was primarily from motor cortices, S2 cortex and thalamus (**Figure 6, Tables 3A, 3B**). In both, Tlx3 and Sim1 brains, long-range input arose from neurons in L2/3 and L5 and double labelled neurons -- i.e., neurons that were presynaptic to L5 cells and projected to L1 of S1 -- were most commonly found in L2/3 (one-way ANOVA, ****p>0.0001). We focused our count of presynaptic neurons on the areas that provide input to L1. In S2 of Tlx3-Cre brains, presynaptic input was from L2/3 and L5a neurons (with a smaller input from L6a and L6b), and 49 neurons were double labelled for fb and mCherry (**Figure 6A**). None of the L6 neurons were double labelled.

**Figure 6.**
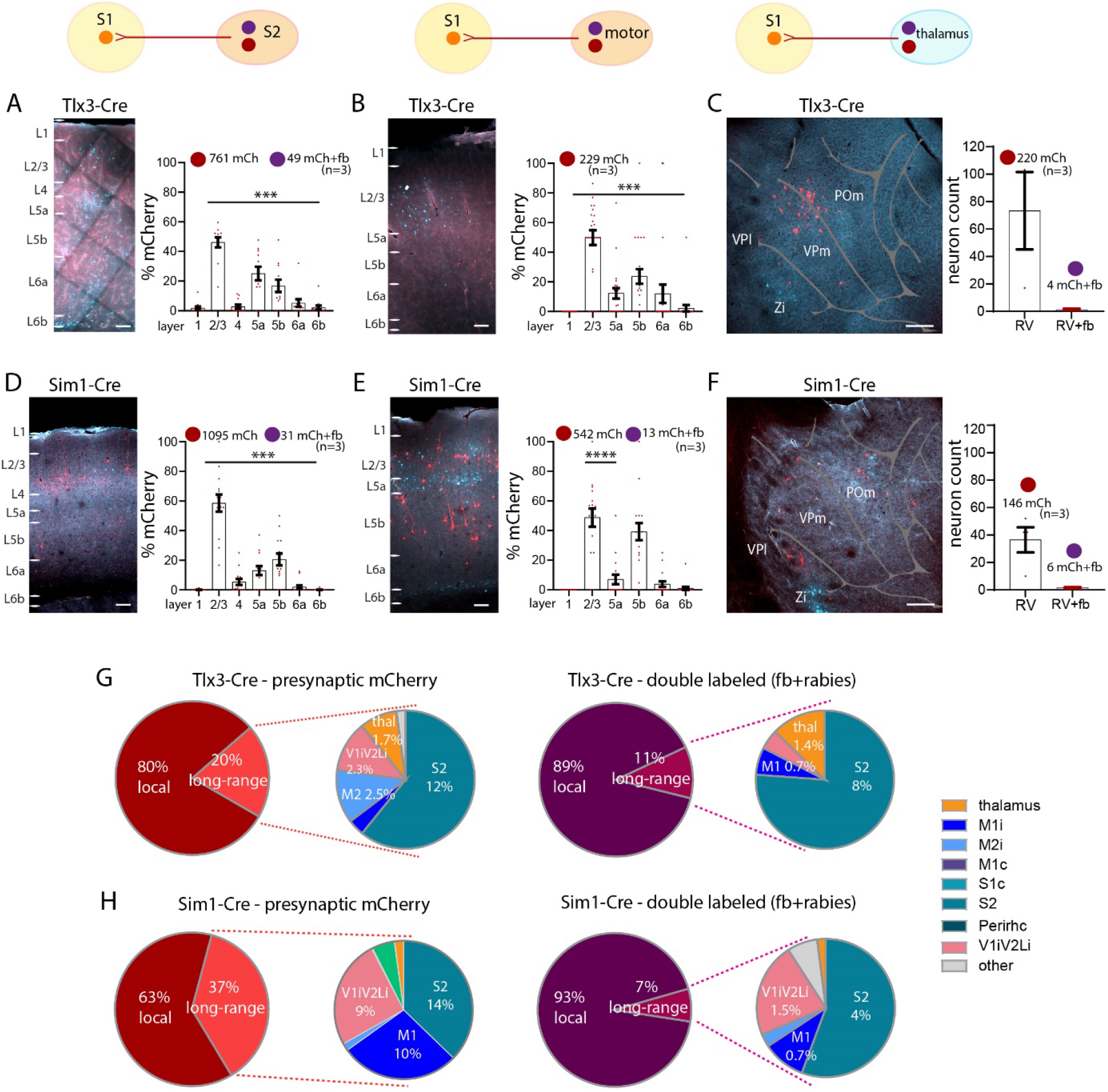
Long-range synaptic input to S1 L1 in Tlx3 and Sim1 neurons in combination with fb. **(A-C)** Images and quantification of presynaptic neurons (rabies and fb double labelled) for Tlx3 brains in **(A)** S2 cortex ipsilateral, in (**B**) motor cortices, and in **(C)** thalamus. **(D-F)** Images and quantification of presynaptic neurons (rabies and fb double labelled) for Sim1 brains, in **(D)** S2 cortex ipsilateral, in (**E**) motor cortices ipsilateral, in (**F**) thalamus. **(G)** Pie charts showing presynaptic input to S1 for both IT neurons (*left*) and L1 projecting neurons that are presynaptic to L5 neurons (*right*). (**H)** Pie charts showing presynaptic input to S1 for both PT neurons (*left*) and L1 projecting neurons that are presynaptic to L5 neurons (*right*). Most input to IT and PT neurons was local. The long-range input to S1 L1 was from S2, Visual (V1 and V2L), M1 and thalamus. For IT neurons the bulk of the neurons presynaptic to these cells that also projected to L1 were local neurons (89%). Presynaptic input to PT neurons in S1 was divided into 63% local and 37% long-range. The long-range input was from S2, Visual (V1 and V2L), and M1. Presynaptic input to PT neurons that also targeted L1 arose from local neurons (93% of the total input). Total number of neurons counted in each mouse line, data from three brains each genotype, shown as mean ± S.E.M. Each dot in the graphs (in **A-F**) represents one brain section. Statistical analysis with one-way ANOVA, Bonferroni post-hoc test, ****p>0.0001. Analysis details in **Tables 3A, 3B, 3D**. Scale bar in **A, B, D, E,** 100 µm, in **C, F**, 500 µm.

L2/3 and L5a neurons in motor cortex provided most of the presynaptic input to S1 Tlx3 neurons, and few were double labelled (**Figure 6B**). Some presynaptic neurons were found in L6a and none were double labelled.

In S2 of Sim1-Cre brains, L2/3 and L5b neurons generated the bulk of the presynaptic input and 31 neurons were double labelled (**Figure 6D**). Presynaptic neurons in motor cortex were found in L2/3, L5a and L5b, but only 13 were double labelled (**Figure 6E**). Presynaptic neurons to both L5 lines were found in thalamus with slightly more input to Tlx3 neurons than to Sim1 neurons. Double labelled neurons accounted for a small percentage of labelled thalamic neurons in both lines (Tlx3: 1.8%, Sim1: 4.1%, **Figure 6C, 6F**).

### Local versus long-range input to L1

Taken together these results suggest that the bulk of the presynaptic input to Tlx3 and Sim1 neurons was from local neurons (Tlx3: 80%, Sim1: 63%). A large portion of the input targeting L1 that was presynaptic to L5 neurons, arose from local neurons. A fraction of the neurons presynaptic to L5 pyramidal neurons also projected to L1. Long-range cortical input to these pyramidal cells was from S2, motor and visual cortices (**Figure 6G, 6H, Table 3D**).

When we extended the approach of counting fb labelled neurons to the whole brain, and to assess the input from the different brain areas, we found that the local input to L1 of S1 arises predominantly from neurons underlying the application site (68%), with long-range input constituting 32% of the total input (n=4, 11956±2469 neurons per brain, **Figure 7A**, **Table 4**). The long-range input to L1 was from higher order thalamus (7%), ipsilateral motor cortices (9%), S2 cortex (2%), visual cortices (3%), and perirhinal cortex (0.1% ipsi and 0.4% contra). Contralateral S1, motor, and perirhinal cortex also contributed input to L1 of somatosensory cortex. We did not quantify input from brainstem to L1. We obtained a similar input pattern when injecting retrobeads in L1: 67% was local, and 33% was long-range input (n=3, 3487±868 neurons per brain, **Figure 7A**, **Table 4**).

**Figure 7.**
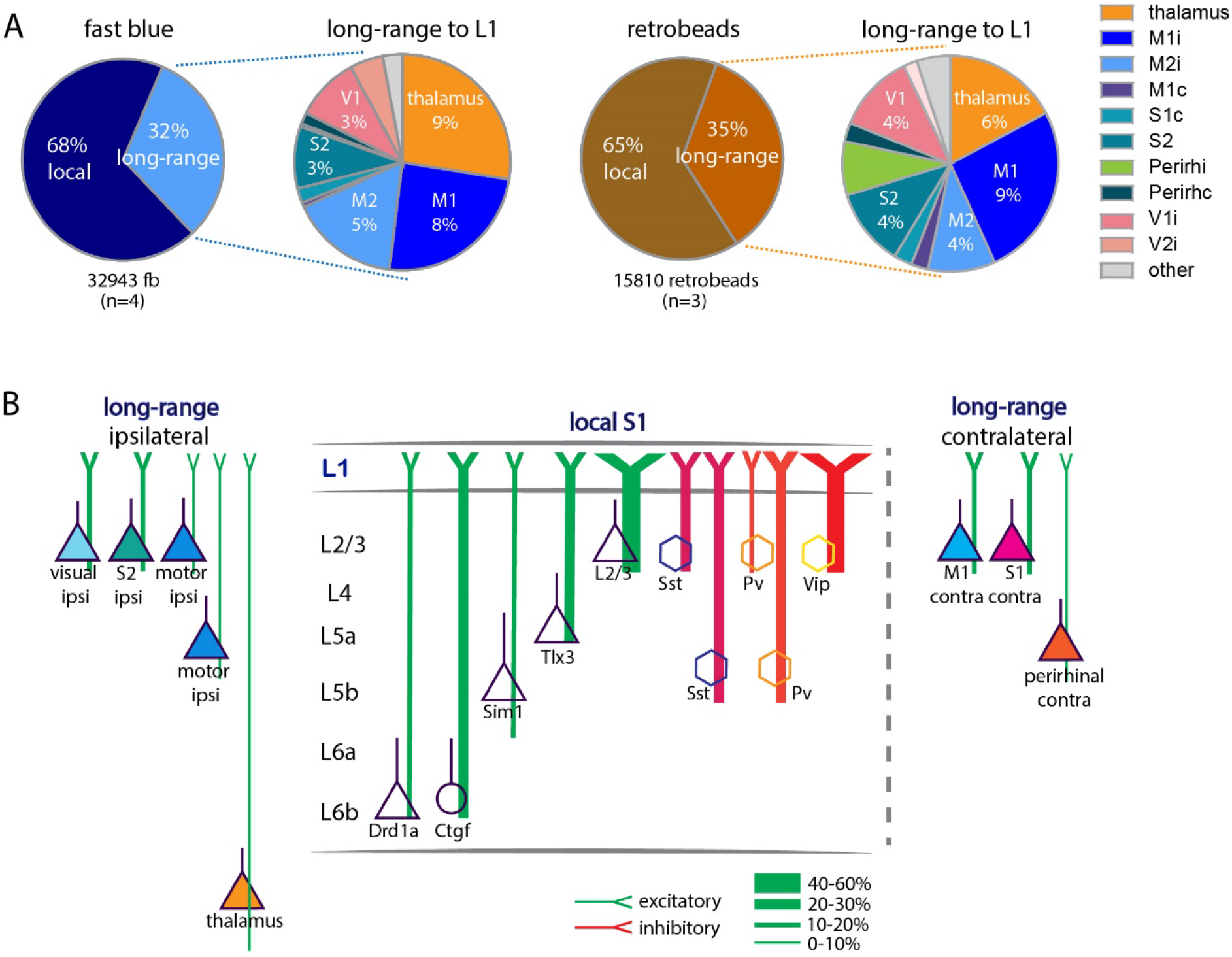
Comparison of anatomical input to L1. **(A)** Schematic of input to L1 assessed with fb *(left)* and retrobeads *(right)*. The bulk of input to L1 was from local neurons and a fraction of the input arising from long-range sources. Thalamus (including POm, zona incerta), ipsilateral and contralateral motor cortices, contralateral S1, ipsilateral S2, bilateral perirhinal (perirh), and ipsilateral visual cortices all provide input to L1. **(B)** Schematic of input to L1. Proportion of local and long-range input and classification of long-range input to L1. Local input arose mainly from L2/3 and L5. Long-range input arose from S2, motor and visual cortices ipsilateral to the application site and was less pronounced from the contralateral S1 and M1. Abbreviations in A: i, ipsi; c, contralateral. Data from four mice for fb, three mice for retrobeads, analysis details in **Table 4**.

## Discussion

Layer 1 has been dubbed an enigma, a ‘crowning mystery’ (Hubel 1982; Rudy et al. 2011) with its anatomy, connectivity, and function still barely charted. By combining traditional retrograde tracing, we quantitatively characterized aspects of organization and function of this layer in primary somatosensory cortex. Our work reveals the extent to which multiple classes of local and long-range neurons connect to L1, and it shows the functional consequences of L5 input to L1.

At the outset of our study, it was known that neurons in L2/3, L5 and L6b provide input to L1 and local inputs are in position to activate L1 interneurons, in fact L2/3 neurons activate these interneurons (Brown and Hestrin 2009; L. Cauller 1995; L. J. Cauller et al. 1998; Clancy and Cauller 1999; Narayanan et al. 2015; Oberlaender et al. 2012; Sakmann 2017; Wozny and Williams 2011; Zolnik et al. 2020). It was also known that long-range inputs from a variety of sources target L1, and that the majority of input to neurogliaform interneurons in L1 arise locally (Abs et al. 2018; L. Cauller 1995; L. J. Cauller et al. 1998; Cohen-Kashi Malina et al. 2021; Doron et al. 2020; M. Larkum 2013; Mease et al. 2016; Oda et al. 2004; Ohno et al. 2012; Sakmann 2017; Veinante and Deschênes 2003; Zagha et al. 2013).

### Input to L1 is primarily local and can modulate activity of L1 interneurons

It is widely accepted that the majority of synapses in any patch of cortex arise from cortical neurons with thalamic inputs, contributing ∼15% of all synapses in cortex, even in L4 (Binzegger et al. 2004; Douglas and Martin 2004; Shepherd 2004; White and Keller 1989). An earlier, detailed analysis of axons and boutons of 39 filled excitatory and inhibitory neurons of all layers of the cat visual cortex, suggested that non-local sources could provide a majority of the input -- "dark synapses" -- to L1 (Binzegger et al. 2004; Boucsein et al. 2011; Douglas and Martin 2004, 2007; Voges et al. 2010). We do not directly address the origin of these mysterious boutons, but along with earlier studies which suggest that the bulk of input to L1 interneurons is from a local origin, our work suggests that local inputs contribute most of the input to L1 (Abs et al. 2018; Brown and Hestrin 2009; Manns et al. 2004; Narayanan et al. 2015; Peng et al. 2021; Sakmann 2017; Wozny and Williams 2011). An additional feature of cortical organization is a substantial recurrent local excitatory and inhibitory connectivity (Sachdev et al. 2012; Sherman and Usrey 2021). L1 neurons are inhibitory and could, via inhibitory interactions in L1 disinhibit pyramidal neurons and interneurons in L2/3 (Anastasiades et al. 2021; Cohen-Kashi Malina et al. 2021).

Our work suggests that local recurrent inputs are important in L1 as well. Neurons with dendrites in L1 -- i.e. L2/3 and L5 pyramidal neurons or L2/3 VIP / SST neurons -- and neurons with axons in L1 contribute local recurrent excitation and inhibition in this layer (M. E. Larkum et al. 2018; Wozny and Williams 2011). Using optogenetic approaches, we show that input to L1 from L5 IT pyramidal neurons has specific functional consequences. L5 IT neurons effectively activate L1 interneurons but do not have a measurable effect on the apical tuft dendrites of L5 PT pyramidal neurons. The negligible effect of L5 IT pyramidal neurons on apical dendrites of L5 PT neurons could reflect weak connectivity between these cell types in L1, or a specific lack of connectivity between the two classes of neurons. L5 pyramidal neurons do not seem to generate local recurrent excitation to other L5 pyramidal neurons via their inputs to L1. It has been shown earlier that uncaging in dendrites can elicit measurable input to the soma (Dodt et al. 1998; Frick et al. 2001; Harris and Pettit 2007; Pettit and Augustine 2000; Shoham et al. 2005), results which are consistent with ours. Taken together our data suggests that circuit mechanisms exist for driving local recurrent excitation and inhibition through synapses in L1 (Abs et al. 2018; Anastasiades et al. 2021; Sakmann 2017; Schuman et al. 2021; Wozny and Williams 2011).

### Long-range input to L1 is primarily from L2/3 and L5 and is cell-type specific

Our work shows that the principal source of long-range input to L1 is from L2/3 neurons, with a smaller but significant input from L5 cortical pyramidal cells. While neurons in contralateral S1, M1 and perirhinal cortices, and ipsilateral S2, M1, M2, perirhinal and visual cortices provide long-range input to L1, the bulk of the long-range input to L1 is from S2, M1, M2, visual cortices and thalamus (**Figure 7B**). Our work also shows that 10% to 20% of the PT and IT neurons in S2 and M1 and contralateral S1 project to L1. While L6 cortico-cortical neurons provide some intra-cortical feedback, their input to L1 is negligible. In contrast to the results with fb applied on the cortical surface, when tracer was injected into S1 cortex, an additional class of cortico-cortical projection neurons in L6 was evident, data which relate to previous studies on hierarchical models with retrograde tracers (Markov et al. 2014; Vezoli et al. 2021).

### Few neurons that are presynaptic to L5 pyramidal neurons project to L1

Previous work has shown that motor cortices, POm thalamus, and perirhinal cortex project to L1 and have substantial input to L5 (Aronoff et al. 2010; Avermann et al. 2012; Chen et al. 2013; Chen et al. 2015; Chen et al. 2016; Doron et al. 2020; Geng et al. 2022; van der Bourg et al. 2017; Yamashita et al. 2018; Zagha et al. 2013). Our experiments with rabies virus shows that input from higher order somatosensory cortex, S2, to L5 PT and IT neurons constitutes more than 40% of the total long-range input to these neurons. Additionally, the rabies approach shows that some L6a and L6b neurons in S2 and M1 are presynaptic to the IT neurons. Nevertheless, even for S2, only a fraction of neurons that were presynaptic to PT and IT neurons also projected to L1. Note that Tlx3 and Sim1 neurons are only subclasses of all L5 neurons, consequently these connectivity numbers are likely to be underestimates.

### Input to L1 in other cortices

The approach we have taken here to study input to L1 in S1 cortex has also been successfully applied to other cortices such as motor/ prefontal cortices (Arbuthnott et al. 1990; Geng et al. 2022; Herkenham 1979). Tracing of axons of filled thalamic neurons confirms that just as in S1 cortex where higher order POm axons target L1 (Sermet et al. 2019; Wimmer et al. 2010), higher order thalamic VM neurons target cortical L1 (Kuramoto et al. 2009; Kuramoto et al. 2015; Kuramoto et al. 2017) and can affect movement initiation (Takahashi et al. 2021).

### Methodological considerations

#### Why use fast blue applied on the cortical surface?

Traditional anatomical approaches for assessing connectivity between cortical areas use retrograde tracers injected into the brain. More recently AAV-retro and pseudo-rabies approaches have been developed targeting Cre-expressing neurons. But input to L1 cannot be assessed with beads, cholera toxin, fast blue or AAV-retro when injected into cortex. The rabies approach is specific and can target specific classes of neurons i.e. L1 interneurons or pyramidal cells with dendrites in L1, but there is no approach that targets pyramidal cell dendrites exclusively. Approaches targeting L1 interneurons can be specific (Abs et al. 2018), but only reveal input to a tiny fraction of all interneurons in cortex and even in L1, only to 60% of interneurons in L1 (Rudy et al. 2011; Schuman et al. 2019).

The use of fb and related tracers applied on the cortical surface was established ∼30 years ago by Cauller and colleagues (L. J. Cauller et al. 1998; Clancy and Cauller 1999; Keizer et al. 1983; Kuypers et al. 1980). This work showed long-range connectivity to L1 and revealed that local L6b and L5 connect to L1 of somatosensory cortex in rats. A concern of retrograde tracing with fb applied on the cortical surface is that fb could spread into L2/3 or even L4. To mitigate against this, we discarded brains in which fb label is found in VPm or VPl. VPm neurons project to L4 and can target the apical tufts specifically in L2/3 (Feldmeyer 2012; Guest et al. 2021; Killackey and Ebner 1973; Oberlaender et al. 2012; Sermet et al. 2019; Wimmer et al. 2010). This approach mitigates against but does not completely eliminate connections that might target L2/3.

Another concern with fb application is the potential for non-specific labelling of neurons directly at the centre of the application site i.e. too much background or potential uptake of fb by dendrites of pyramidal neurons. To address this issue we used four approaches: 1) We injected a different class of retrograde tracers -- retrobeads -- and compared the results we obtained with retrobeads to those with fb. When we used retrobeads we observe sparser label locally and in distant cortical and subcortical structures, but the overall pattern of label with both retrobeads and fb are similar; 2) We considered the functional consequences of local input to L1. Earlier physiology with single cell fills of pyramidal neurons indicated that L2/3 axons were pervasive in L1 and generated substantial input to L1 (Wozny and Williams 2011). As with L2/3, single cell fills revealed that L5a neurons have dense projections to L1, but while L5b axons are present in L1, the local axon is sparser (Sakmann 2017). In our study we tested the physiological effect of these local axons on both L1 interneurons and L5 pyramidal cells and show a specific functional effect of L5a input to L1 interneurons. The interaction between L5 cells, their input to L5 dendrites, in L1 was negligible; 3) We counted both at centre of the application site where there was dense fb label, where it was possible to overestimate labelled neurons, and 140 microns off centre of the application site. The overall pattern of label was similar in both cases; 4) We injected fb into cortex and found a different laminar pattern of both local and long-range label.

One additional concern with all tracing approaches is the uptake of tracer by fibers of passage. While this is possible, it should be noted that all tracing methods seem to show a similar pattern of labelling, with neurons in the same cortical and subcortical areas labelled. Our labelling with fb is consistent from mice to mice, generating a similar laminar profile for inputs. Uptake by fibres of passage might have generated a less consistent pattern of label.

#### Definition of cortical layers

While L1 and L6b are easily defined, the borders that separate the other layers from each other are not easily identifiable. We defined layers using 100/200-micron bins. This approach has the benefit of consistency and reproducibility, but there can be errors arising from brain shrinkage, brain size or selection of sections used for counting. To mitigate against this, we examined the proportion of fb labelled neurons in a 30-micron strip at the centre of the 100/200-micron bin defining each layer. The proportion of fb labelled neurons in L4 changed with this approach, suggesting that some of the L4 label could be attributed to L2/3 or L5.

#### Rabies virus tracing

The incubation time we used for rabies tracing (seven days) might have been too short to obtain expression in all long-range neurons (Kim et al., 2016). Furthermore, even though we have ruled out leakage of rabies virus and found little non-specific expression (Zolnik et al. 2020), the rabies virus approach could be marginally non-specific.

### Implication

The nub of the mystery concerning L1 can be reduced to two questions: 1) How is the input to this layer organized, and 2) What does the input do? Our work shows that L1 has some features that are common to neocortical organization. Most input to this layer is local, arising from excitatory and inhibitory neurons in the patch of cortex underlying L1. This implies that when feedforward, feedback, or both inputs drive somatic action potentials in L2/3 or L5, this activity is likely to directly generate synaptic activity in L1, thus likely to modulate the effect of a backpropagating action potentials that interact with contextual long-range input.

Our characterization of anatomical and functional input to L1 in S1 cortex adds to the growing knowledge and reveals key organizational principles for this circuit. Our work also points to a need to develop tools to further understanding cortical organization and function. Overall, our work suggests that local input to L1 may play a critical role in the information flowing through the underlying cortex.

## Acknowledgements

We would like to thank: The Viral Core Facility of the Charité – Universitätsmedizin Berlin for their support and the generation of the viruses used in this study; the AMBIO Facility of the Charité – Universitätsmedizin Berlin for their support; Kristin Lehmann for excellent technical assistance; Justus Donner for help with tissue processing; Albert Gidon for input on an earlier version of the manuscript; Benjamin Judkewitz for providing the Drd1a-Cre and Ctgf-dgCre lines; Jörg Geiger for providing the Pv-IRES-Cre-Ai9 line; Ehud Ahissar for the Gpr26-Cre line; Michael Brecht for providing the Scnn1A-Cre line; Charles Gerfen for Tlx3-Cre and Sim1-Cre lines; James Poulet for importing the Gpr26-Cre line and sharing the Vip-IRES-Cre line; and Edward Callaway for the following generous gifts: pAAV-EF1a-DIOHTB, pAAV-Ef1a-DIO-H2B-GFP-2A-oG-WPRE-hGH, cDNA B7GG, BHKEnvA cells, and HEK293T-TVA cells.

## Author contributions

J.M.T.L., T.A.Z., R.N.S., and M.E.L. planned and designed the experiments and wrote the manuscript. J.L., T.A.Z., and R.N.S. performed the experiments. J.M.T.L., T.A.Z., R.N.S., and M.T. analyzed the data. T.T., and C.R. produced viruses, designed experiments and wrote the manuscript. B.J.E. and D.J. wrote the manuscript. All authors approved the final version of the manuscript.

## Funding

This work was supported by the European Union’s Horizon 2020 research and innovation program and Euratom research and training program 20142018 (under grant agreement No. 670118 to MEL); Deutsche Forschungsgemeinschaft (Exc 257 NeuroCure, Grant No. LA 3442/3-1 & Grant No. LA, Project number 327654276 SFB1315); European Union Horizon 2020 Research and Innovation Program (72070/HBP SGA1, 785907/HBP SGA2, 785907/HBP SGA3 670118/ERC Active Cortex); Einstein Foundation Berlin (EVF-2017-363), and NINDS (R01NS1114702).

## Conflict of interests

The authors declare no competing financial interests.

## Notes

### Competing Interest Statement

The authors have declared no competing interest.

### Summary of Updates

We have improved the manuscript, text and images according to suggestions of three reviewers.

