## Supplementary figures and tables for "Layer 1 of somatosensory cortex: An important site for input to a tiny cortical compartment"

1, Institute of Biology, Humboldt Universität zu Berlin Charitéplatz 1, Virchowweg 6, 10117 Berlin, Germany; 2, Institute of Neurophysiology, Charité – Universitätsmedizin Berlin, Charitéplatz 1, Virchowweg 6, 10117 Berlin, Germany; 3, Institute of Biochemistry, Charité – Universitätsmedizin Berlin, Charitéplatz 1, Virchowweg 6, 10117 Berlin, Germany; 4, Neurocure Centre for Excellence Charité – Universitätsmedizin Berlin; 5, Emory University, Atlanta, GA, USA.

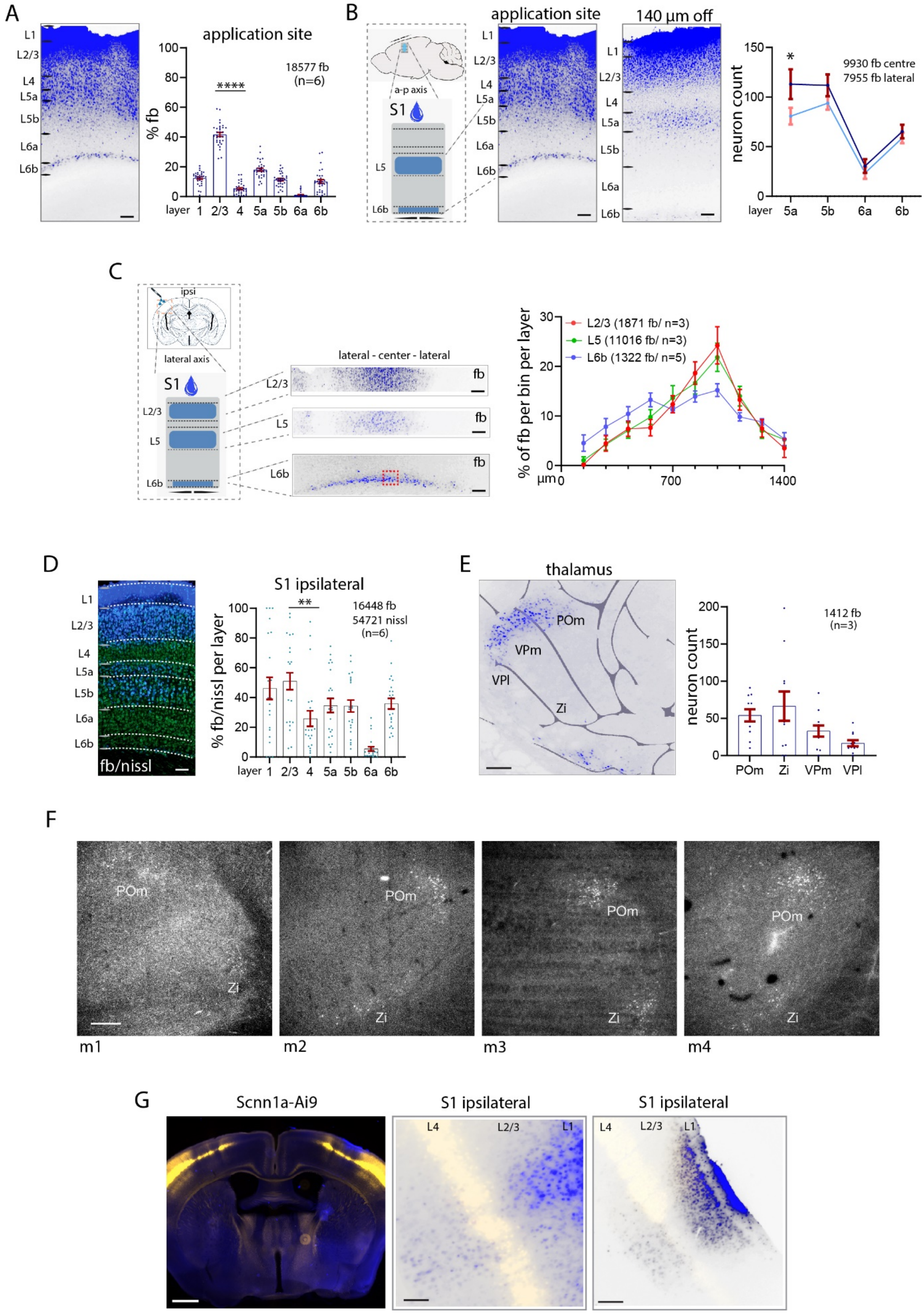
A
application site
18577 fb
(n=6)
% fb
layer
B
a-p axis
S1
application site
140 µm off
9930 fb centre
7955 fb lateral
neuron count
C
ipsi
lateral axis
lateral - center - lateral
fb
L2/3 (1871 fb/ n=3)
L5 (11016 fb/ n=3)
L6b (1322 fb/ n=5)
% of fb per bin per layer
µm
D
fb/nissl
S1 ipsilateral
16448 fb
54721 nissl
(n=6)
% fb/nissl per layer
E
thalamus
POm
VPm
VPl
Zi
1412 fb
(n=3)
F
m1
m2
m3
m4
G
Scnn1a-Ai9
S1 ipsilateral
S1 ipsilateral

**Supplementary Figure 1. Local input to S1 L1.** (**A)** Uptake of fb at the application site in S1 cortex including center brain sections. **(B)** Fewer fb neurons were labelled in L5 and L6 (7955 fb, 31 brain sections) compared to the application site (9930 fb, 31 brain sections, one-way ANOVA *$*p<0.05$*). (**C**) In lateral direction, fb label diminished laterally towards the edges from the center in L6b. The distribution of fb label in L2/3 and L5 is not significantly narrower than in L6b (750 fb neurons). (**D**) Laminar percentages for fb uptake at the application site in 100-micron bins. **(E)** Brains with fb label in VPl and VPm were excluded from analysis. Total numbers of neurons counts are shown in each panel. Statistical analysis with one-way ANOVA, Bonferroni post-hoc test, ****$p<0.0001$. (**F**) Example images showing fb label in POm (**G**) Example images of fb label in Scnn1a-Cre-Ai9 brain sections showing that fb label has not been taken up by cells in L4**.** Each dot in the graphs represents one brain section. Data from mice as indicated (in brackets). Analysis details in **Table 1A**. Scale bars in **A, B, D,** 100 µm, in **C**, and **G** *(right)* 50 µm, in **E**, **F**, and **G** 500 µm.

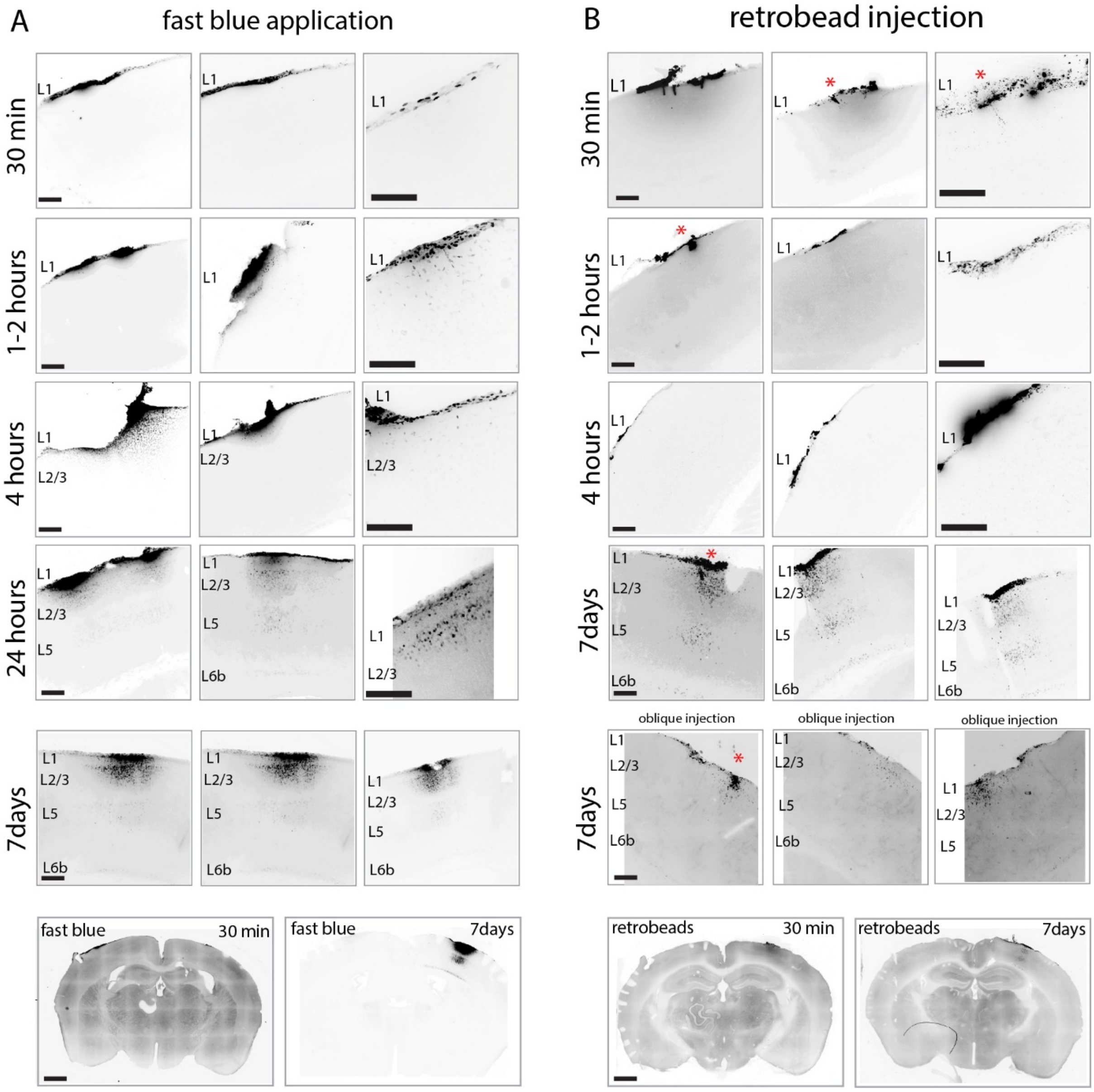


**Supplementary Figure 2. Application of fast blue and injection of retrobeads in a time series. (A)** Application of fb after ½ hour, 1-2 hours, 4 hours, 24 hours and 7 days showing uptake of fast blue in deeper layers 2/3 around 4 hours. **(B)** Injection of retrobeads after ½ hour, 1-2 hours, 4 hours, and 7 days (with oblique injection method) showing uptake of red fluorescence in deeper layers after 4 hours. Red asterisk marking injection site. Scale bars in **A, B**, 100 µm, in higher magnification images 50 µm, in overview images 500 µm.

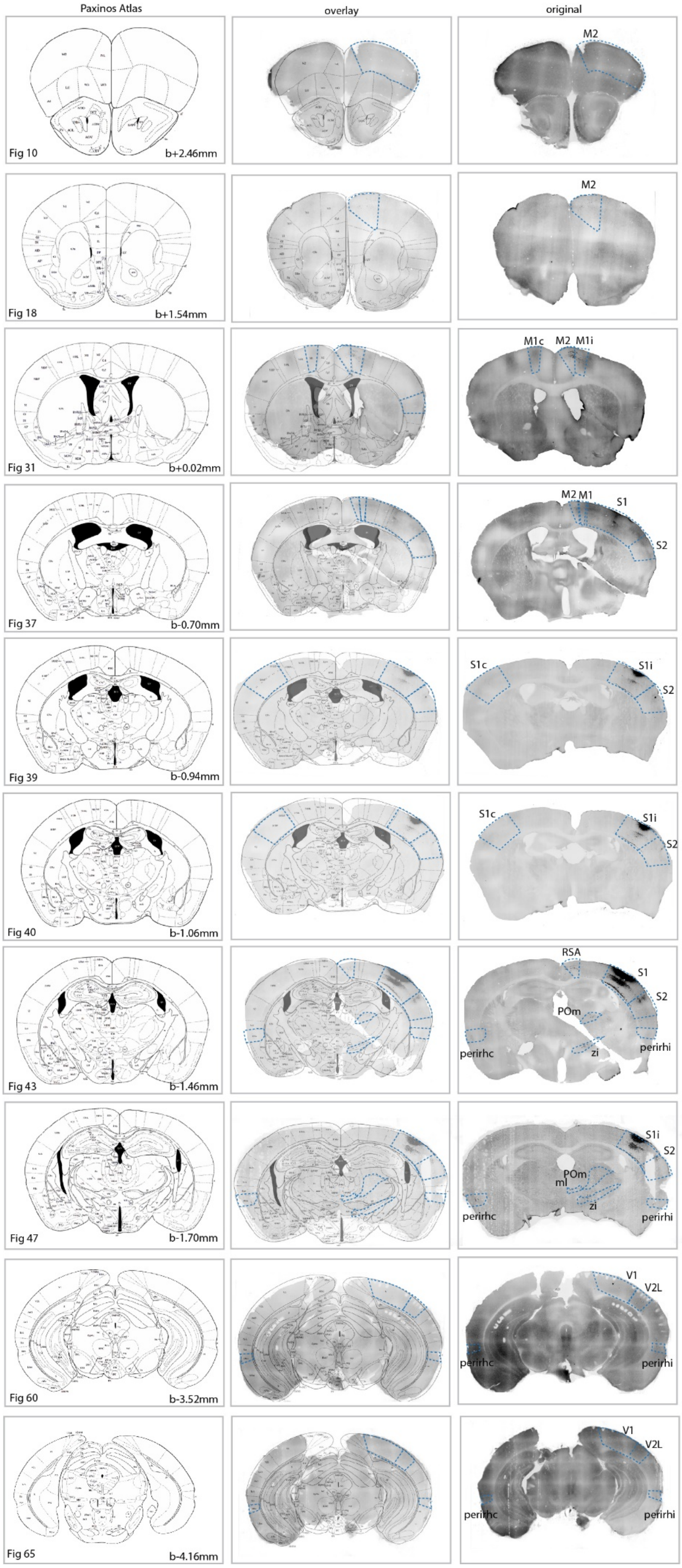
Paxinos Atlas
overlay
original
M2
Fig 10
b+2.46mm
M2
Fig 18
b+1.54mm
M1c
M2
M1i
Fig 31
b+0.02mm
M2
M1
S1
S2
Fig 37
b-0.70mm
S1c
S1i
S2
Fig 39
b-0.94mm
S1c
S1i
S2
Fig 40
b-1.06mm
RSA
S1
S2
POm
perirhc
zi
perirhi
Fig 43
b-1.46mm
S1i
S2
POm
ml
perirhc
zi
perirhi
Fig 47
b-1.70mm
V1
V2L
perirhc
perirhi
Fig 60
b-3.52mm
V1
V2L
perirhc
perirhi
Fig 65
b-4.16mm

**Supplementary Figure 3. Fast blue images aligned to the reference atlas.** Images were aligned to the Paxinos reference atlas. Images from anterior-posterior direction showing fb label in distinct brain areas. Areas with fb label are marked with blue outlines and indicated as motor cortices M1, M2, somatosensory cortices S1, S2, (i, ipsi, c, contra), visual cortices V1, V2L, perirhinal cortex, perirhi (ipsi), perirhc (contra), POm thalamus, Zi zona incerta.

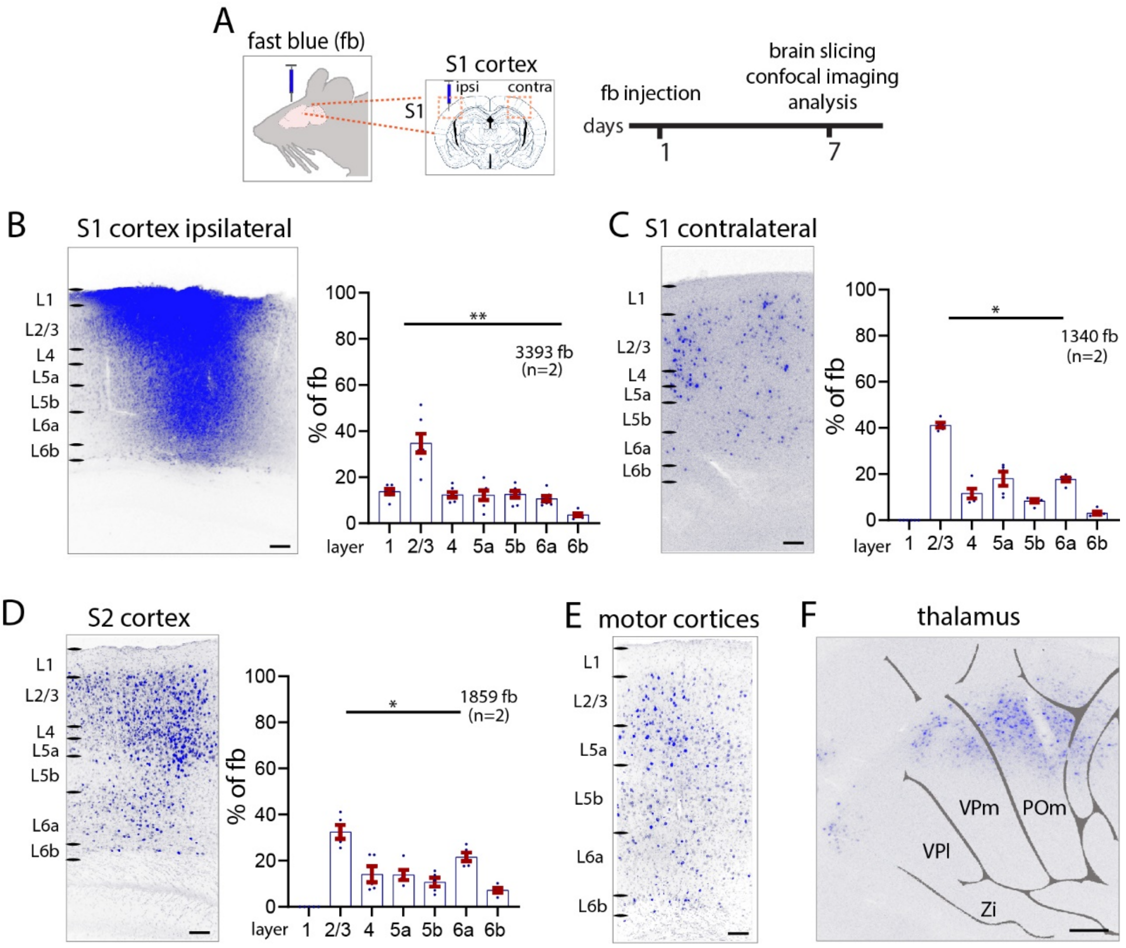


**Supplementary Figure 4. Fast blue injection in S1 cortex.** (**A**) Schematic for injection of fb into S1 cortex. **(B)** When fb was injected into cortex, fb labelled neurons were distributed in a different pattern than when applied on cortex L1. Fast blue labelled neurons were found in all cortical layers, primarily in L2/3. (**C**) Example images and quantification of fb labelling in contralateral S1, and (**D**) in ipsilateral S2 cortex. (**E**) Example images of motor cortex show fb labelled neurons in all layers. (**F**) Example images of fb uptake in POm, VPm, and VPl thalamus. Each dot in the graphs represents one brain section. Total number of neurons counted and mice used (in brackets) are shown in each panel. Data from two mice. One-way ANOVA, Bonferroni post-hoc test, **p<0.01. Analysis details in **Table 1C.** Scale bars in **B-E**, 100 µm, in **F** 500 µm.

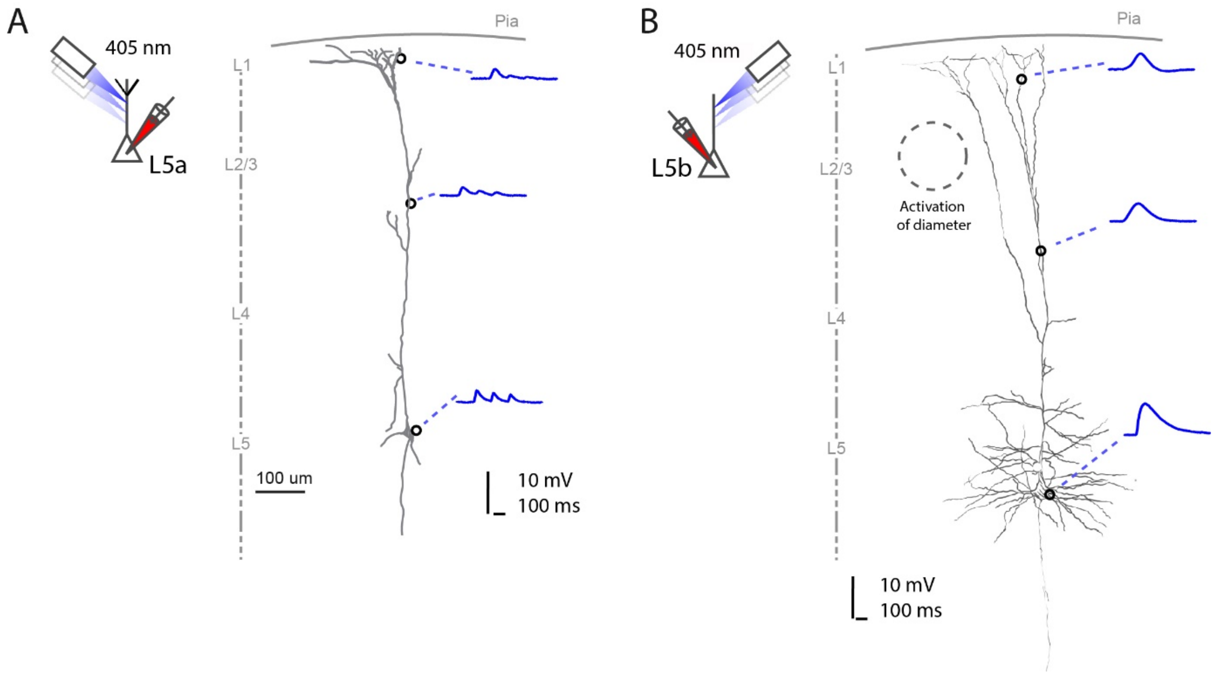


**Supplementary Figure 5. Glutamate uncaging in L5a and L5b neurons revealed EPSPs along the dendrites.** Experimental setup to test input along the L5a **(A)** and L5b **(B)** neuron. EPSP traces in reconstructed neurons after glutamate uncaging was performed at the soma, at the apical dendrites, and at the apical tufts in L1 (n=5 neurons, three mice used per genotype).

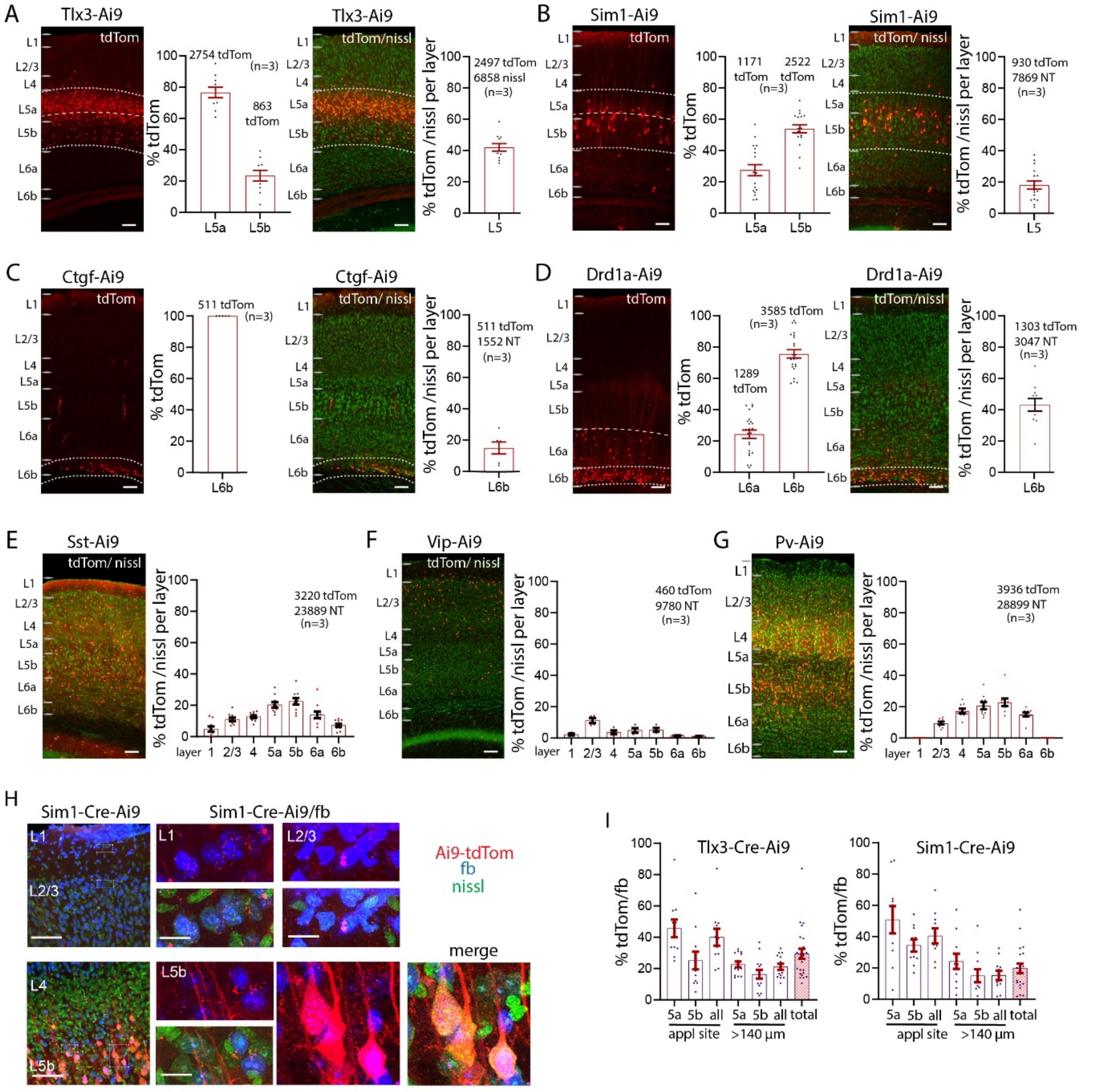

**Supplementary Figure 6. Local input from classes of L5 and L6b, and inhibitory cells to L1. (A)** Images of sections showing IT Tlx3-Cre/ tdTom positive and the nissl stained neurons and quantification. Seventy-six percent of tdTom positive cells in this line were in L5a. Forty-two percent of all L5 cells were tdTom positive. (**B**) Sections showing PT Sim1-Cre/ tdTom positive and nissl stained neurons and quantification. In this line, 18% of L5 neurons were tdTom positive. (**C**) Proportion of L6b neurons labelled in Ctgf-Cre/ tdTom positive and nissl stained neurons. All Ctgf neurons are located in L6b. Fifteen percent of tdTom positive cells in this line were in L6b. **(D)** In the Drd1a-Cre line 76% of tdTom positive cells are located in L6b, the other remaining in L6a. Forty-two percent of all L6b neurons in L6b is Drd1a positive. **(E)** Pattern of Sst/ tdTom double labelled neurons. Quantification of laminar pattern of Sst positive neurons with nissl stain. (**F**) Pattern of Vip/ tdTom double labelled neurons. Quantification of laminar pattern shows percentages of Vip neurons. (**G**) Pattern of Pv/ tdTom double labelled neurons. Quantification of laminar pattern shows percentages of Pv neurons. **(H)** High magnification images of fb double labelled Sim1-Cre-Ai9 neurons show no uptake of fb in dendrites. **(I)** Quantification of fb uptake in Tlx3-Cre-Ai9 and in Sim1-Cre-Ai9 in L5a, L5b, and total L5, at the centre and >140 microns distant from the application site. Analysis details see **Table 2E.** Each dot in the graphs represents one brain section. Total number of neurons counted and mice (in brackets) used are shown in each panel. Data from three mice each genotype. Analysis details in **Tables 2A, 2B, 2E**. Fast blue pseudo colored in cyan. Scale bars 100 µm.

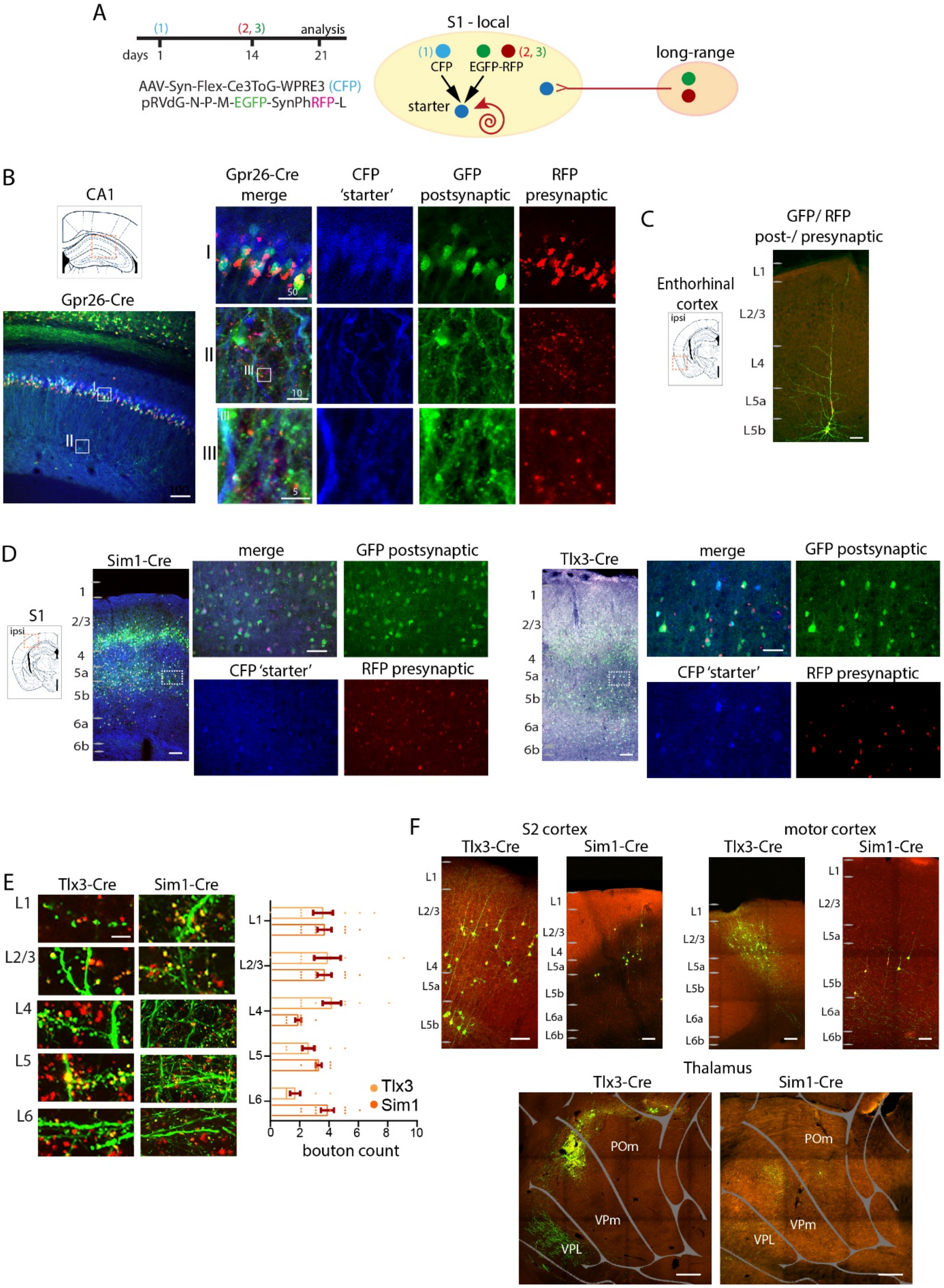
A
days 1 14 21
(1) (2, 3) analysis
AAV-Syn-Flex-Ce3ToG-WPRE3 (CFP)
pRVdG-N-P-M-EGFP-SynPhRFP-L
S1 - local
(1) CFP
EGFP-RFP (2, 3)
starter
long-range
B
CA1
Gpr26-Cre
Gpr26-Cre merge
CFP 'starter'
GFP postsynaptic
RFP presynaptic
I
II
III
C
GFP/ RFP post-/ presynaptic
Enthorhinal cortex
ipsi
L1
L2/3
L4
L5a
L5b
D
S1
ipsi
Sim1-Cre
merge
GFP postsynaptic
CFP 'starter'
RFP presynaptic
Tlx3-Cre
E
Tlx3-Cre
Sim1-Cre
L1
L2/3
L4
L5
L6
bouton count
Tlx3
Sim1
F
S2 cortex
motor cortex
Tlx3-Cre
Sim1-Cre
Thalamus
POm
VPm
VPL

**Supplementary Figure 7. Synaptic input to S1 L1. (A)** Schematic showing injection scheme for rabies virus tracing in Gpr26-Cre, Tlx3-Cre and Sim1-Cre mice. (**B**) Example images of CA1 hippocampus in Gpr26-Cre mice, with CFP positive starter neurons, GFP positive postsynaptic neurons, RFP positive presynaptic boutons and merged images. (**C**) Example image of entorhinal cortex confirming long-range projections to CA1. (**D**) Example images of Tlx3-Cre and Sim1-Cre brains of S1 cortex with CFP positive starter neurons, GFP positive postsynaptic neurons, RFP positive presynaptic boutons and merged images. (**E**) Quantification of boutons (yellow) in Tlx3-Cre and Sim1-Cre brains across the layers (Tlx3, n=369 synapses, Sim1, n=158 synapses, counting from 10 boxes in each layer with size of 20 µm x 10 µm, 1 dot represents 1 box). (**F**) Example images in S2, motor cortex and thalamus showing presynaptic neurons. Three brains per genotype, one-way ANOVA, Bonferroni post-hoc test, ***$p<0.001$. Data shown as mean ± S.E.M. Data from three mice each genotype. Scale bars in **B-D, F**, 100 µm, in zoom-ins as indicated or 50 µm, in **E** 5 µm, in **F** thalamus 500 µm.

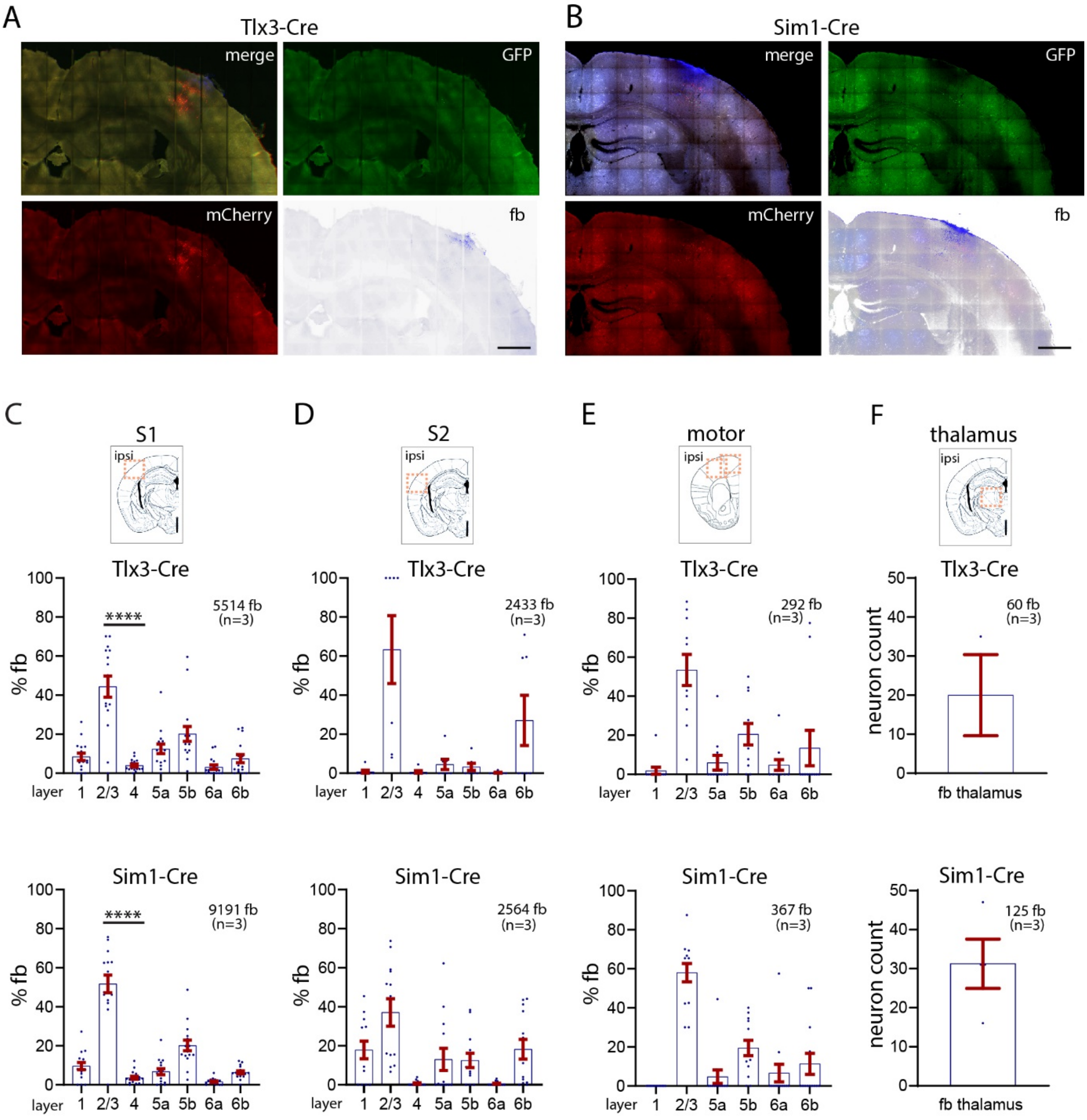


**Supplementary Figure 8. S1 injection sites and fb labelled neurons in rabies experiments.** (**A, B**) Example images of injection sites of rabies virus in Tlx3-Cre and Sim1-Cre brains. (**C-F**) Quantification of fb labelled neurons in Tlx3-Cre and Sim1-Cre brains of rabies tracing. (**C**) In S1 cortex ipsilateral, (**D**) S2 cortex ipsilateral, (**E**) Motor cortices ipsilateral, (**F**) Thalamus (neuron counts). Data shown as mean ± S.E.M. Each dot in the graphs represents one brain section. Total number of neurons counted and mice used (in brackets) are shown in each panel.

Data from three mice each genotype. Statistical analysis with one-way ANOVA, Bonferroni post-hoc test, ****p<0.0001. Analysis details in **Tables 3A, 3B**. Scale bars in **A, B**, 800 μm.

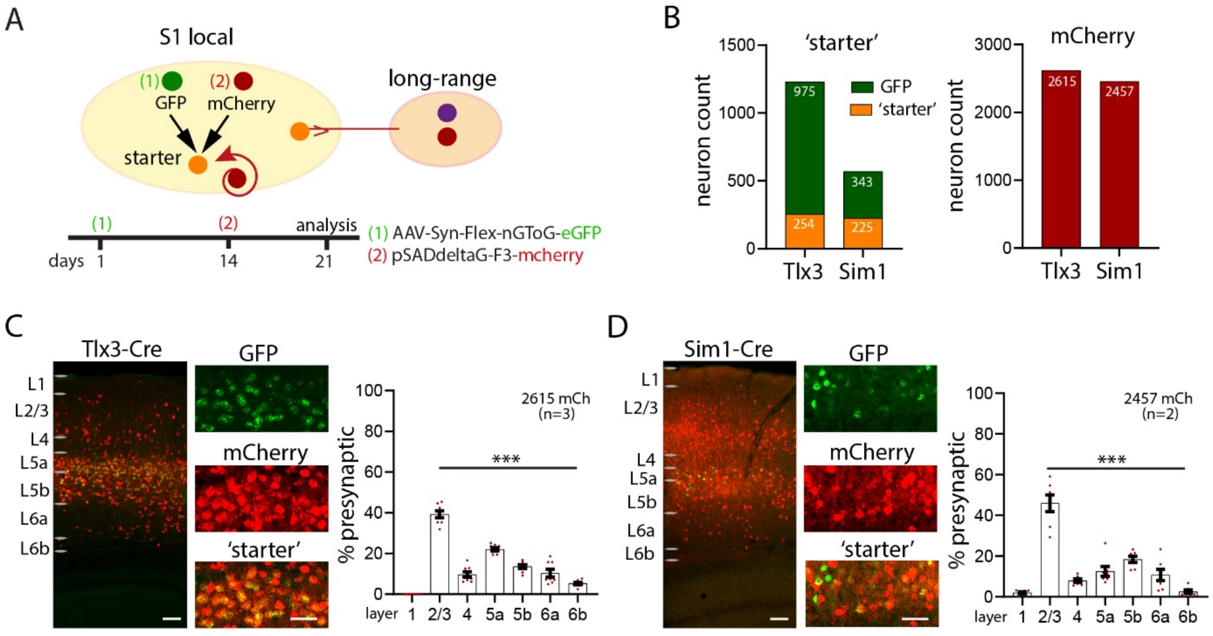


**Supplementary Figure 9. Rabies virus in Tlx3-Cre and Sim1-Cre brains.** (**A**) Schematic showing timeline for synaptic rabies virus tracing without fb application. (**B**) In Tlx3-Cre, the presynaptic input (2615 mCherry) was derived from 254 starter neurons (975 GFP neurons). In Sim1-Cre, the presynaptic input (2457 mCherry) was derived from 225 starter neurons (343 GFP neurons). (**C, D**) Example images of Tlx3-Cre brains and Sim1-Cre brains in S1 cortex at the injection sites. Higher magnification shows starter, GFP, and mCherry neurons. Quantification of presynaptic neurons in Tlx3-Cre and Sim1-Cre brains in graphs. Each dot represents one brain section. Total number of neurons counted and mice used (in brackets) are shown in each panel. One-way ANOVA, Bonferroni post-hoc test, ***p<0.001. Data shown as mean ± S.E.M. Analysis details in **Table 3C.** Scale bars in **C, D** 100 μm, in zoom-ins 50 μm.

### Supplementary Tables

#### Table 1A: Input to L1 with fast blue application and retrobead injection.

| % fb, cortices (number of neurons, average per brain) | L1 | L2/3 | L4 | L5a | L5b | L6a | L6b | Mice/ brain sections |
|---|---|---|---|---|---|---|---|---|
| **fb S1 ipsi without center sections (**9868 fb) | 12.7±1.3 | 39.8±1.9 **** | 4.8±0.9 | 17.9±1.4 | 10.5±1.2 | 1.4±0.5 | 12.8±1.9 | 6/13 (**Figure 1B**) |
| **fb S1 ipsi all sections** (18577 fb, 3096±459) | 12.4±0.8 | 41.7±1.4 **** | 5.3±0.7 | 18.1±1.0 | 11.4±0.8 | 1.0±0.3 | 10.0±1.3 | 6/31 **(Suppl Figure 1A**) |
| **fb/nissl S1 ipsi, 30-micron bin** (1071 fb, 3664 nissl) | 42.7±15.7 | 45.2±11.9 ** | 4.2±0.8 | 32.9±8.8 | 20.4±6.3 | 0.4±0.3 | 31.9±5.5 | 3/8 (**Figure 1C**) |
| **fb/nissl S1 ipsi (**16448 fb, 54721 nissl) | 46.1±7.4 | 51.0±5.8 **** | 25.8±5.1 | 34.6±4.7 | 34.3±4.0 | 5.4±1.4 | 35.9±3.5 | 6/34 **(Suppl Figure 1D**) |
| **retrobeads** (5391 retrobeads, 2490±374) | 15.3±2.0 | 44.9±2.4 **** | 0.1±0.09 | 11.6±1.0 | 11.3±0.9 | 0.3±0.3 | 16.4±2.1 | 3/30 (**Figure 1D**) |

Values are percentage mean ± standard error, data from mice as indicated (**Figure 1, Supplementary Figure 1**). Statistical analysis one-way ANOVA, ****p<0.0001. P-values indicate significance level for comparison between L2/3 and other layers. Comparison between layers in fast blue application and retrobead injections; L6a** (Kruskal-Wallis test, p<0.0001).

#### Table 1B: Input from other cortical areas to L1 S1 cortex.

| % fb, cortices (number of neurons, average per brain) | L1 | L2/3 | L4 | L5a | L5b | L6a | L6b |
|---|---|---|---|---|---|---|---|
| **fb S1 contra (**1883 fb, 366±55) | 0±0 | 61.0±2.9 **** | 1.7±0.5 | 24.9±2.3 | 12.0±1.7 | 0.3±0.2 | 0.03±0.03 |
| **fb/nissl S1 contra (**1221 fb, 102, 4239 nissl) | 0±0 | 9.4±8.0 **** | 0±0 | 4.0±1.5 | 2.9±1.2 | 0±0 | 0±0 |
| **fb M2 ipsi** (1311 fb) | 0.1±0.1 | 36.6±5.2 | - | 48.4±5.7 | 9.9±2.0 | 4.6±1.1 | 0.4±0.4 |
| **fb/nissl M2 ipsi** (1311 fb, 13327 nissl) | 0.2±0.1 | 14.2±3.1 | - | 34.7±6.0 **** | 6.6±1.5 | 2.3±0.8 | 0.1±0.1 |
| **fb M1 ipsi** (1343 fb) | 0±0 | 50.2±6.2 | - | 31.1±7.1 | 9.4±2.1 | 2.9±1.3 | 6.5±5.4 |
| **fb/nissl M1 ipsi** (1343 fb, 12285 nissl) | 0±0 | 19.4±3.0 | - | 20.8±5.4 | 5.4±1.3 | 1.9±0.9 | 3.6±2.9 |

| | | | | | | | |
|---|---|---|---|---|---|---|---|
| **fb S2 ipsi** (1663 fb) | 0±0 | 41.8±6.2 | 8.8±1.2 | 12.9±1.6 | 30.4±10.7 | 10.1±5.3 | 56.0±1.7 |
| **fb/nissl S2 ipsi** (1663 fb, 11763 nissl) | 0±0 | 42.3±14.0 | 15.9±4.1 **** | 14.0±1.1 | 8.6±2.0 | 10.7±2.1 | 20.1±2.2 |
| **fb visual ipsi** (1056 fb) | 1.6±1.6 | 50.7±2.4 | 5.7±14.8 | 14.9±2.5 | 9.7±3.9 | 2.0±1.4 | 15.4±7.4 |
| **fb/nissl visual ipsi** (1056 fb, 5544 nissl) | 6.1±6.1 | 41.5±14.2 | 18.5±15.0 **** | 23.3±10.2 | 14.4±4.5 | 4.1±3.5 | 11.1±2.4 |
| **fb perih ipsi** (51 fb) | 0±0 | 1.6±1.6 | 0±0 | 90.3±5.6 | 8.1±4.9 | 0±0 | 0±0 |
| **fb perih contra** (146 fb) | 0±0 | 7.2±4.2 | 0±0 | 70.5±7.4 | 22.4±8.6 | 0±0 | 0±0 |

Values are percentage mean ± standard error, data from four mice (**Figure 2**). Statistical analysis one-way ANOVA, ****p<0.0001. P-values indicate significance level for comparison between L2/3 (or L5a, M2) and other layers.

#### Table 1C: Fast blue injection into S1 cortex.

| % fb, cortices/ layers, (number of neurons) | L1 | L2/3 | L4 | L5a | L5b | L6a | L6b | Mice/ brain sections |
|---|---|---|---|---|---|---|---|---|
| **fb injection S1 ipsi** (3393 fb) | 13.8±1.1 | 34.8±4.1 ** | 12.5±1.2 | 12.2±2.0 | 12.6±1.4 | 10.7±1.2 | 3.7±0.6 | 2/5 (**Suppl Figure 4B**) |
| **fb injection S1 contra** (1340 fb) | 0±0 | 41.3±1.1 * | 11.6±2.1 | 18.0±3.0 | 17.7±1.0 | 3.0±0.7 | 0±0 | 2/5 (**Suppl Figure 4C**) |
| **fb injection S2 ipsi** (1859 fb) | 0±0 | 32.5±3.0 * | 14.2±3.5 | 13.8±2.2 | 10.7±1.9 | 21.7±1.9 | 7.2±1.0 | 2/5 (**Suppl Figure 4D**) |

Values are percentage mean ± standard error, data from two mice (**Supplementary Figure 4**). Statistical analysis one-way ANOVA, ****p<0.0001. P-values indicate significance level for comparison between L2/3 and other layers. Comparison between layers in fast blue application, retrobead injections, and fast blue injection; L4****, L6a**** (Kruskal-Wallis test, p<0.0001).

#### Table 2A: tdTom expression in excitatory Ai9-Cre lines.

| % fb, cortices/ layers, (number of neurons) | L5a | L5b | L6b | Mice/ brain sections |
|---|---|---|---|---|
| **Tlx3-Ai9, S1 ipsi** (3617 tdTom) | 75.6±2.4 | 23.8±2.3 | - | 3/16 (**Suppl Figure 6A**) |
| **Tlx3-Ai9, S1 ipsi, nissl** (2497 tdTom, 6858 nissl) | 42.1±2.4 | - | - | 3/11 (**Suppl Figure 6A**) |
| **Sim1-Ai9, S1 ipsi** (3693 tdTom) | 27.5±3.6 | 54.0±2.5 | - | 3/18 (**Suppl Figure 6B**) |
| **Sim1-Ai9, S1 ipsi, nissl** (930 tdTom, 7869 nissl) | - | 18.1±2.6 | - | 3/16 (**Suppl Figure 6B**) |
| **Ctgf-Ai9, S1 ipsi, nissl** (1552 tdTom, 511 nissl) | - | - | 14.9±3.8 | 3/6 (**Suppl Figure 6C**) |
| **Drd1a-Ai9, S1 ipsi** (4874 tdTom) | - | - | 75.7±2.6 | 4/23 (**Suppl Figure 6D**) |
| **Drd1a-Ai9, S1 ipsi, nissl** (1303 tdTom, 3047 nissl) | - | - | 42.8±4.0 | 3/11 (**Suppl Figure 6D**) |

Values are percentage mean ± standard error, data from three mice each genotype and as indicated (**Supplementary Figure 6**).

#### Table 2B: tdTom expression in inhibitory Ai9-Cre lines.

| Cortices / layers, Tlx3 (%) | L 1 | L 2/3 | L4 | L5a | L5b | L6a | L6b | Mice/ brain sections |
|---|---|---|---|---|---|---|---|---|
| **Sst-Ai9, S1 ipsi, nissl** (3220 tdTom, 23889 nissl) | 4.4±1.6 | 11.0±1.1 | 12.7±0.8 | 20.3±1.7 | 22.5±1.7 | 13.9±2.2 | 7.3±1.0 | 3/10 (**Suppl Figure 6E**) |
| **Vip-Ai9, S1 ipsi, nissl** (460 tdTom, 9780 nissl) | 2.3±0.5 | 11.3±1.6 | 3.6±1.2 | 4.9±1.4 | 5.2±1.2 | 1.5±0.2 | 1.2±0.2 | 4/10 (**Suppl Figure 6F**) |
| **PV-Ai9, S1 ipsi, nissl** (3936 tdTom, 28899 nissl) | 0±0 | 9.6±0.9 | 17.3±1.5 | 20.7±2.3 | 22.9±2.4 | 15.1±1.3 | 0±0 | 3/9 (**Suppl Figure 6G**) |

Values are percentage mean ± standard error, data from three mice each genotype and as indicated (**Supplementary Figure 6**).

#### Table 2C: Input from classes of L5 and L6b cells to L1.

| % fb, cortices/ layers, (number of neurons) | L5a | L5b | L6b | Mice/ brain sections |
|---|---|---|---|---|

| | | | | |
|---|---|---|---|---|
| **Tlx3-Ai9 S1 ipsi, tdTom/fb** (7706 tdTom, 2361 tdTom+fb) | 29.4±3.2 **** | - | - | 3/25 (**Figure 4A**) |
| **Sim1-Ai9, S1 ipsi, tdTom/fb** (1832 tdTom, 382 tdTom+fb) | - | 19.7±3.0 | - | 3/22 (**Figure 4B**) |
| **Ctgf-Ai9, S1 ipsi, fb** (578 tdTom, 288 tdTom+fb) | - | - | 51.5±9. 9 | 3/12 (**Figure 4C**) |
| **Drd1a-Ai9, S1 ipsi, fb** (3447 tdTom, 510 tdTom+fb) | - | - | 12.7±7.0 | 3/18 (**Figure 4D**) |
| **Tlx3-Ai9, S1 contra, tdTom/fb** (2100 tdTom, 111 tdTom+fb) | 7.7±1.4 | - | - | 3/18 (**Figure 4H**) |
| **Tlx3-Ai9, S2 ipsi, tdTom/fb** (439 tdTom, 49 tdTom+fb) | 11.1±1.1 | - | - | 3/3 (**Figure 4H**) |
| **Tlx3-Ai9, motor ipsi, tdTom/fb** (1249 tdTom, 141 tdTom+fb) | 10.7±1.7 | - | - | 3/15 (**Figure 4H**) |
| **Sim1-Ai9, S1 contra, tdTom/fb** (4522 tdTom, 73 tdTom+fb) | - | 1.3±0.5 | - | 3/18 (**Figure 4I**) |
| **Sim1-Ai9, S2 ipsi, tdTom/fb** (412 tdTom, 19 tdTom+fb) | - | 8.2±1.5 | - | 3/13 (**Figure 4I**) |
| **Sim1-Ai9, motor ipsi, tdTom/fb** (1083 tdTom, 42 tdTom+fb) | - | 10.6±2.1 | - | 3/12 (**Figure 4I**) |

Values are percentage mean ± standard error, data from three mice each genotype (**Figure 4**).

#### Table 2D: Input from inhibitory SST, VIP, and PV cells to L1.

| **% fb, cortices/ layers, (number of neurons)** | **L 1** | **L 2/3** | **L4** | **L5a** | **L5b** | **L6a** | **L6b** | **Mice/ brain sections** |
|---|---|---|---|---|---|---|---|---|
| **Sst-Ai9, S1 ipsi, tdTom/fb** (3819 tdTom, 377 tdTom+fb) | 28.3±7.2 | 19.7±2.3 | 12.1±2.2 | 11.6±1.5 | 8.9±1.1 | 1.5±0.6 | 3.0±1.0 | 3/25 (**Figure 4E**) |
| **Vip-Ai9, S1 ipsi, tdTom/fb** (777 tdTom, 93 tdTom+fb) | 4.0±2.5 | 81.2±4.9 **** | 2.0±1.2 | 5.3±2.4 | 5.9±2.7 | 0±0 | 1.6±1.6 | 3/7 (**Figure 4F**) |
| **PV-Ai9, S1 ipsi, tdTom/fb** (3942 tdTom, 318 tdTom+fb) | 0±0 | 14.2±2.7 | 0±0 | 15.9±3.3 | 19.7±3.5 | 0±0 | 0±0 | 3/15 (**Figure 4G**) |

Values are percentage mean ± standard error, data from three mice each genotype (**Figure 4**).

#### Table 2E: Fb application - Tlx3-Cre-Ai9 and Sim1-Cre-Ai9 in L5.

| **% fb, cortices/ layers, (number of neurons)** | **L5a** | **L5b** | **L5** | **sections** |
|---|---|---|---|---|

| | | | | |
|---|---|---|---|---|
| **Tlx3-Ai9, S1i tdTom/fb** (5746 tdTom, 1575 tdTom +fb) | 45.66±6.60 (c), 22.64±1.82 (l) | 34.34±8.68 (c), 14.98±4.17 (l) | 40.03±5.44 (c), 21.08±1.99 (l) | 25 **(Suppl Figure 6)** |
| **Sim1-Ai9, S1i, tdTom/fb** (4038 tdTom, 1209 tdTom +fb) | 50.74±8.68 (c), 24.16±4.90 (l) | 25.15±5.65 (c), 16.13±2.78 (l) | 40.43±4.81 (c), 15.22±3.04 (l) | 22 **(Suppl Figure 6)** |

Values are percentage mean ± standard error, data from three mice each genotype (**Supplementary Figure 6**).

#### Table 3A: Rabies virus and fast blue application in Tlx3-Cre mice.

| **Cortices / layers, Tlx3 (%)** | **L 1** | **L 2/3** | **L4** | **L5a** | **L5b** | **L6a** | **L6b** | **sections** |
|---|---|---|---|---|---|---|---|---|
| **mCherry S1** (6138 total, 4728 local) | 3.6±1.4 | 37.8±2.8 **** | 8.9±0.9 | 23.7±1.8 | 16.8±1.8 | 7.9±1.6 | 1.2±1.1 | 15 (**Figure 5C**) |
| **fb S1** (9596 total, 5514 local) | 9.7±1.9 | 47.1±4.8 **** | 3.3±0.8 | 14.6±1.0 | 13.4±3.5 | 1.4±0.6 | 10.4±2.9 | 10 **(Suppl Figure 8C)** |
| **fb-mCherry S1** (449 total, 391 local) | 4.78±1.4 | 26.8±3.78 **** | 9.43±1.9 | 28.1±3.1 | 30.9±3.17 | 0±0 | 0±0 | 15 (**Figure 5C**) |
| **mCherry S2** (761) | 1.79±3.65 | 46.2±3.38 *** | 2.80±1.31 | 25.1±4.57 | 16.8±4.16 | 5.22±2.53 | 2.13±1.37 | 9 (**Figure 6A**) |
| **fb S2** (2433) | 0.8±0.8 | 63.3±17.4 | 0.6±0.6 | 4.5±2.7 | 3.3±1.9 | 0.2±0.2 | 27.1±12.9 | **9 (Suppl Figure 8D)** |
| **mCherry motor** (229) | 0±0 | 49.9±4.99 *** | - | 12.3±3.56 | 23.7±4.94 | 12.0±6.20 | 2.17±2.17 | 12 (**Figure 6B**) |
| **fb motor** (292) | 1.8±1.8 | 53.5±26.4 | - | 5.9±3.7 | 20.5±5.5 | 4.8±2.7 | 13.5±9.0 | 11 **(Suppl Figure 8E)** |

Values are percentage mean ± standard error, data from three mice **(Figures 5, 6, Supplementary Figure 8**). Statistical analysis one-way ANOVA, ****$p<0.0001$. P-values indicate significance level for comparison between L2/3 and other layers.

#### Table 3B: Rabies virus and fast blue application in Sim1-Cre mice.

| **Cortices / layers, Sim1 (%)** | **L 1** | **L 2/3** | **L4** | **L5a** | **L5b** | **L6a** | **L6b** | **sections** |
|---|---|---|---|---|---|---|---|---|
| **mCherry S1** (6495 total, 3869 local) | 1.5±0.2 | 38.6±1.8 **** | 6.6±1.0 | 8.5±0.7 | 36.7±1.33 **** | 6.94±1.1 | 1.18±0.18 | 15 (**Figure 5D**) |
| **fb S1** (13131 total, 9191 local) | 13.7±3.9 | 43.0±4.8 **** | 1.2±0.4 | 26.0±4.7 | 3.8±1.3 | 0.6±0.3 | 11.6±2.8 | 12 **(Suppl Figure 8C)** |
| **fb-mCherry S1** (823 total, 754 local) | 3.9±0.9 | 38.1±5.6 **** | 2.4±1.1 | 7.6±1.9 | 46.6±5.5 **** | 0.3±0.3 | 0.3±0.2 | 15 (**Figure 5D**) |

| | | | | | | | | |
|---|---|---|---|---|---|---|---|---|
| **mCherry S2** (1095) | 0.1±0.1 | 58.6±5.8 *** | 5.4±2.01 | 13.1±3.0 | 20.6±4.0 | 1.9±1.2 | 0.1±0.13 | 14 (**Figure 6D**) |
| **fb S2** (2564) | 17.9±4.5 | 37.1±7.0 | 0.6±0.4 | 13.0±5.6 | 12.5±3.6 | 0.6±0.3 | 18.2±5.10 | 14 **(Suppl Figure 8D)** |
| **fb-mCherry S2** (31) | 0±0 | 19.0±7.7 | 0±0 | 66.2±14.5 | 14.8±8.4 | 0±0 | 0±0 | 6 (**Figure 6D**) |
| **mCherry motor** (542) | 0±0 | 48.8±6.1 **** | - | 7.1±3.3 | 39.2±5.7 | 3.9±1.9 | 1.0±1.0 | 17 (**Figure 6E**) |
| **fb motor** (367) | 0±0 | 58.0±4.7 | - | 4.7±3.4 | 19.4±3.9 | 6.5±4.5 | 11.4±5.4 | 13 **(Suppl Figure 8E)** |
| **fb-mCherry motor** (13) | 0±0 | 30.7±11.9 | - | 1.8±1.8 | 3.2±2.2 | 0±0 | 0±0 | 9 (**Figure 6E**) |

Values are percentage mean ± standard error, data from three mice (**Figures 5, 6, Supplementary Figure 8**). Statistical analysis one-way ANOVA, ****p<0.0001. P-values indicate significance level for comparison between L2/3 (L5b) and other layers.

#### Table 3C: Rabies virus in Tlx3-Cre and Sim1-Cre brains.

| **mCherry, layers S1 (%)** | **L 1** | **L 2/3** | **L4** | **L5a** | **L5b** | **L6a** | **L6b** | **sections** |
|---|---|---|---|---|---|---|---|---|
| **Tlx3-Cre** (2615 mCherry) | 0±0 | 39.3±1.7 *** | 9.6±1.4 | 22.1±0.8 | 13.5±0.9 | 10.3±1.9 | 5.2±0.5 | 10 |
| **Sim1-Cre** (2457 mCherry) | 1.9 ±0.5 | 45.9±4.1 *** | 7.9±0.8 | 12.5±2.4 | 18.4±1.5 | 10.7±2.7 | 2.5±0.8 | 9 |

Values are percentage mean ± standard error, data from three mice (Tlx3-Cre), two mice (Sim1-Cre) (**Supplementary Figure 9**). Statistical analysis one-way ANOVA, ***p<0.001. P-values indicate significance level for comparison between L2/3 and other layers.

#### Table 3D: Rabies virus and fast blue application in Tlx3-Cre and Sim1-Cre mice.

| **% (number of neurons)** | **Local** | **Long-range** | **S1c** | **M1i** | **M2i** | **M1c** | **S2i** | **Perirhc** | **V1i, V2Li** | **Thal** | **other** |
|---|---|---|---|---|---|---|---|---|---|---|---|
| **Tlx3 mCherry** | 80.3±5.3 | 19.7±5.3 | 0±0 | 0.8±0.3 | 2.5±2.5 | 0±0 | 12.0±2.4 | 0.01±0.01 | 2.3±1.2 | 1.7±0.9 | 0.4±0.4 |
| **Tlx3 mCherry+fb** | 89.1±7.7 | 10.9±7.7 | 0±0 | 0.7±0.4 | 0±0 | 0±0 | 8.3±8.3 | 0±0 | 0.5±0.5 | 1.3±0.7 | 0±0 |

| | | | | | | | | | | | |
|---|---|---|---|---|---|---|---|---|---|---|---|
| **Sim1 mCherry** | 62.8±5.5 | 37.2±5.5 | 0.1±0.1 | 10.3±5.0 | 0.6±0.4 | 1.9±1.9 | 13.8±5.1 | 0.4±0.4 | 9.5±2.9 | 0.8±0.04 | 0±0 |
| **Sim1 mCherry +fb** | 93.1±2.0 | 6.9±2.0 | 0±0 | 0.7±0.7 | 0.2±0.2 | 0±0 | 3.8±0.1 | 0±0 | 1.5±1.5 | 0.1±0.1 | 0±0 |

Values are percentage mean ± standard error, data from three mice each genotype (**Figure 6**).

#### Table 4: Fast blue application and retrobead injection on S1

| % (number of neurons, average per brain) | Local | Long-range | S1c | M1i | M2i | M1c | S2i | Perirh-i | Perirh-c | V1i | V2Li | Thal | other |
|---|---|---|---|---|---|---|---|---|---|---|---|---|---|
| **fb** (32943, 11956±2469) | 68.0±5.6 | 31.8±5.6 | 0.7±0.5 | 7.7±4.8 | 5.2±3.2 | 0.2±0.2 | 2.9±1.9 | 0.2±0.2 | 0.5±0.5 | 1.6±1.1 | 1.6±1.4 | 8.7±2.6 | 0.9±0.1 |
| **retrobead** (15810, 3487±868) | 64.7±5.3 | 35.3±5.3 | 0.9±0.5 | 9.2±1.6 | 3.6±1.6 | 0.9±0.5 | 4.1±1.0 | 2.8±0.9 | 0.9±0.1 | 4.2±2.7 | 0.7±0.4 | 6.1±2.0 | 1.8±1.6 |

Values are percentage mean ± standard error, data from four mice (fb), three mice (retrobeads) (**Figure 7**).
